# Profiling telomere content and composition at single-cell resolution in cancer and immune cells

**DOI:** 10.1101/2024.08.28.609339

**Authors:** Niklas L. Engel, Hanna Frieß, Julia Grigorjew, Ferdinand Popp, Lea Herzel, Binnur Özay, Julie Surmely, Malte Simon, Charles D. Imbusch, Benedikt Brors, Lars Feuerbach

## Abstract

Biological processes such as aging, carcinogenesis, and immune responses depend on the ability of cells to maintain or rapidly expand populations. This capacity is constrained by a cell’s replicative potential, which is reflected in its telomere content. Despite the central role of telomeres in cancer and immunity, their analysis at single-cell resolution across diverse cell types remains challenging. Here we show that scATAC-seq data enables quantitative telomeromics when key technical and biological confounders are accounted for. We present a computational framework that addresses read sparsity, telomere representation, and chromatin-state-dependent competition for sequencing signal, enabling robust estimation of telomere content and telomeric variant repeat composition from scATAC-seq data. By inferring global chromatin condensation directly from scATAC-seq profiles, our approach corrects for cell-cycle-associated biases while simultaneously capturing chromatin-state dysregulation in cancer. Applied to a large cancer atlas, this framework reveals patient-specific telomere maintenance phenotypes, including telomerase-associated and alternative lengthening of telomeres (ALT)-like profiles preserved across subclonal populations. Extending beyond cancer cells, we observe that telomere content in exhausted T cell subpopulations prior to immunotherapy is predictive for effective response to PD-1 checkpoint blockade. Together, these results establish scATAC-seq as a robust platform for single-cell telomeromics in cancer and immunity.

**Graphical Abstract:** 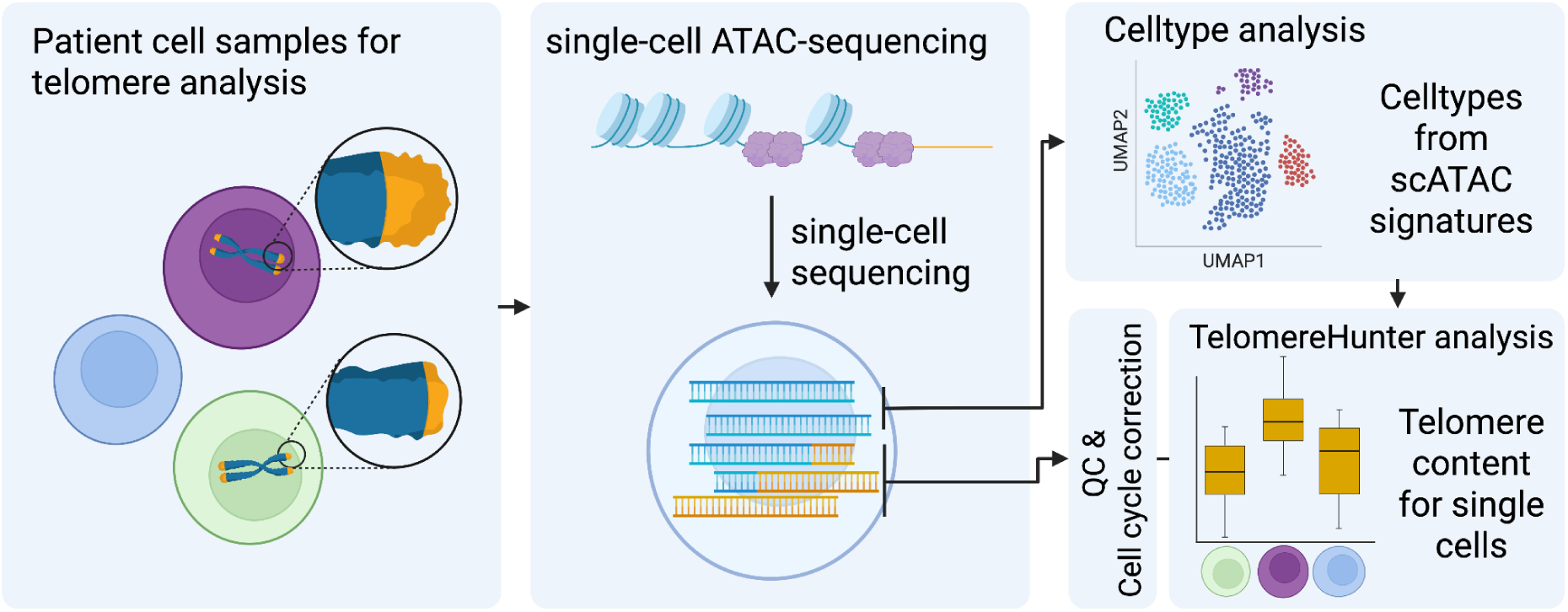

Created in BioRender. Popp, F. (2026) https://BioRender.com/mi2jove

## Introduction

The ends of human chromosomes are protected by telomeres from degradation and fusion with other DNA molecules^1^. Constituted primarily of the hexameric repeat TTAGGG^2,3^, the telomeres form a loop structure stabilized by the proteins of the shelterin complex^4^. Telomeres have a heterogeneous length, which is anti-correlated with age and subject to inherited predisposition^5,6^. Telomeres shorten during cell division due to the end-replication problem^7^, which eventually triggers cellular senescence upon reaching a critical telomere length (M1 checkpoint / Hayflick limit)^8^. Telomere maintenance (TM) counteracts this process, but is not active in all cells. During embryogenesis, the enzyme telomerase elongates telomeres by using the Telomerase RNA component (TERC) as a template to add new nucleotides to telomeric repeats^9^. Later, the expression of the gene encoding the enzymes catalytic subunit, *TERT*, is silenced between the 12th and 18th week of gestation^10,11^. Subsequently, most somatic cells display no telomerase activity. Exceptions include germline cells, stem cells, and components of the adaptive immune system^10,12,13^. Nonetheless, TM is incomplete. More specifically, median telomere length in lymphocytes decreases from ∼10 kbp at birth to ∼4.5 kbp in centenarians^14^, which contributes to aging and loss of immune fitness, and can impact the success of cancer immunotherapy^15^.

In cancer the length and composition of telomeres of the tumor, as well as, the immune cells impact the etiology of the disease. More specifically, the maintenance of telomere length is essential for cellular immortalization and sustained proliferation. While the majority of human cancers achieve this through telomerase reactivation, a substantial minority employ the alternative lengthening of telomeres (ALT) pathway^12,16^. ALT is a telomerase-independent mechanism that relies on homology-directed recombination to elongate telomeric DNA, often resulting in highly heterogeneous telomere lengths and interspersion of telomere variant repeats (TVRs)^17–22^. Comprehensive pan-cancer studies have delineated the genomic footprints of ALT activity across tumor types. ALT signatures, which include TVRs, are enriched in certain sarcomas, gliomas, and pancreatic neuroendocrine tumors^21,23^. More specifically, the hexameres TCAGGG, TGAGGG, TTGGGG, and TTCGGG are enriched in ALT-positive cancers^22^. In contrast, TTTGGG frequency is positively correlated with telomerase activation^24^. These observations suggest that telomere architecture and repeat composition can serve as molecular markers to distinguish ALT-positive from telomerase-positive cancers and provide insights into tumor evolution and genome stability mechanisms.

The adaptive immune system, as the natural antagonist of cancer, consists of a variety of cell types. Among them are T cells, which can be subdivided into CD4⁺ and CD8⁺ T cells. The CD4⁺ subpopulation comprises T helper (Th) and regulatory T cells (Tregs), including T follicular helper cells (Tfh). The CD8⁺ compartment includes effector and memory T cells. CD4⁺ T cells primarily function as helpers, while CD8⁺ T cells have a cytotoxic role. Naïve T cells await activation by their cognate antigen, which triggers differentiation and rapid clonal expansion^25,26^. In cancer, continuous stimulation through TCR and other signals can lead to dysfunctional, exhausted T cells (TEx)^27^, which can be categorized into early, intermediate, and terminal states of exhaustion^28–30^. To sustain proliferation, some T cells expressing the co-receptor CD28 upregulate telomerase via the NF-κB signaling pathway upon activation^31^. Failure of effective immune responses in elderly patients may be linked to the limited replicative potential of lymphocytes due to short telomeres, a process also observed in accelerated form during chronic viral infections^32^.

Single-cell Assay for Transposase-Accessible Chromatin with sequencing (scATAC-seq) is a widely applied protocol that enriches DNA libraries for open chromatin regions. We here describe a computational workflow that solves three central challenges of scATAC-seq-based single-cell telomeromics: read sparsity, telomere representation, and chromatin-state-dependent competition for sequencing signal. Although ATAC-seq is primarily used to identify regions of open chromatin, telomeric reads have been observed to be enriched in such datasets both in cells with intact shelterin and in shelterin-deficient systems^33^. Our study clarifies which coverage per cell is sufficient (read sparsity), and whether shelterin-bound DNA allows unbiased Tn5 transposase activity or if telomeric sequences are as underrepresented as heterochromatin in the resulting datasets (telomere representation). We furthermore develop a correction procedure for the influence of changes in chromatin condensation during the cell cycle on *in silico* telomere quantification in scATAC-seq data, as this process has previously been described as confounder^34^ (competition for sequencing signal).

To this end, we adapted the TelomereHunter software^35^, previously applied to characterize the telomere compartment, respectively telomereome, of whole-genome, exome, and ChIP-seq datasets^22,35–37^. As proof-of-concept, we then applied this approach to a large-scale scATAC-seq dataset recently generated by The Cancer Genome Atlas (TCGA)^38^, comprising multiple cancer types, to investigate telomere composition, heterogeneity, and the distribution of telomere maintenance mechanisms at single-cell resolution across various tumors. We further extended this analysis to a scATAC-seq dataset derived from tumor biopsies of basal cell carcinoma (BCC) patients treated with PD-1 checkpoint blockade^28^, characterizing telomere content differences between immune cell subsets. In this dataset, telomere content of intermediate and terminal exhausted T cells prior to therapy onset predicts therapeutic response, thus underscoring the potential of telomere content as a biomarker for immunotherapy success. We anticipate that the application of the presented scATAC-seq telomeromics workflow on existing and upcoming datasets will provide novel insights into telomere biology of tumor and immune cells, which in turn may have a profound impact on the diagnosis, prognosis and treatment of cancer.

## Results

To perform *in silico* telomeromics based on scATAC-seq data three challenges have to be addressed. (1) Sparseness: Which cells have sufficient sequencing depth to be informative on the telomere content? (2) Representation: Does the ATAC-seq protocol sufficiently represent the telomeres, or does shelterin repel the Tn5 transposase? (3) Competing signals: Do global changes in the chromatin landscape (chromatin plasticity), i.e. during the cell cycle, compete so strongly for the sequencing capacity that telomere content signals are irrevocably confounded? In the following, we address each of these questions using two datasets: The TCGA single-cell atlas dataset from Sundaram *et al.* (2024), a large-scale resource comprising multiple cancer types, and the BCC dataset from Satpathy *et al.* (2019), a scATAC-seq dataset derived from tumor biopsies of BCC patients treated with PD-1 checkpoint blockade (Fig. 1).

**Figure 1.**
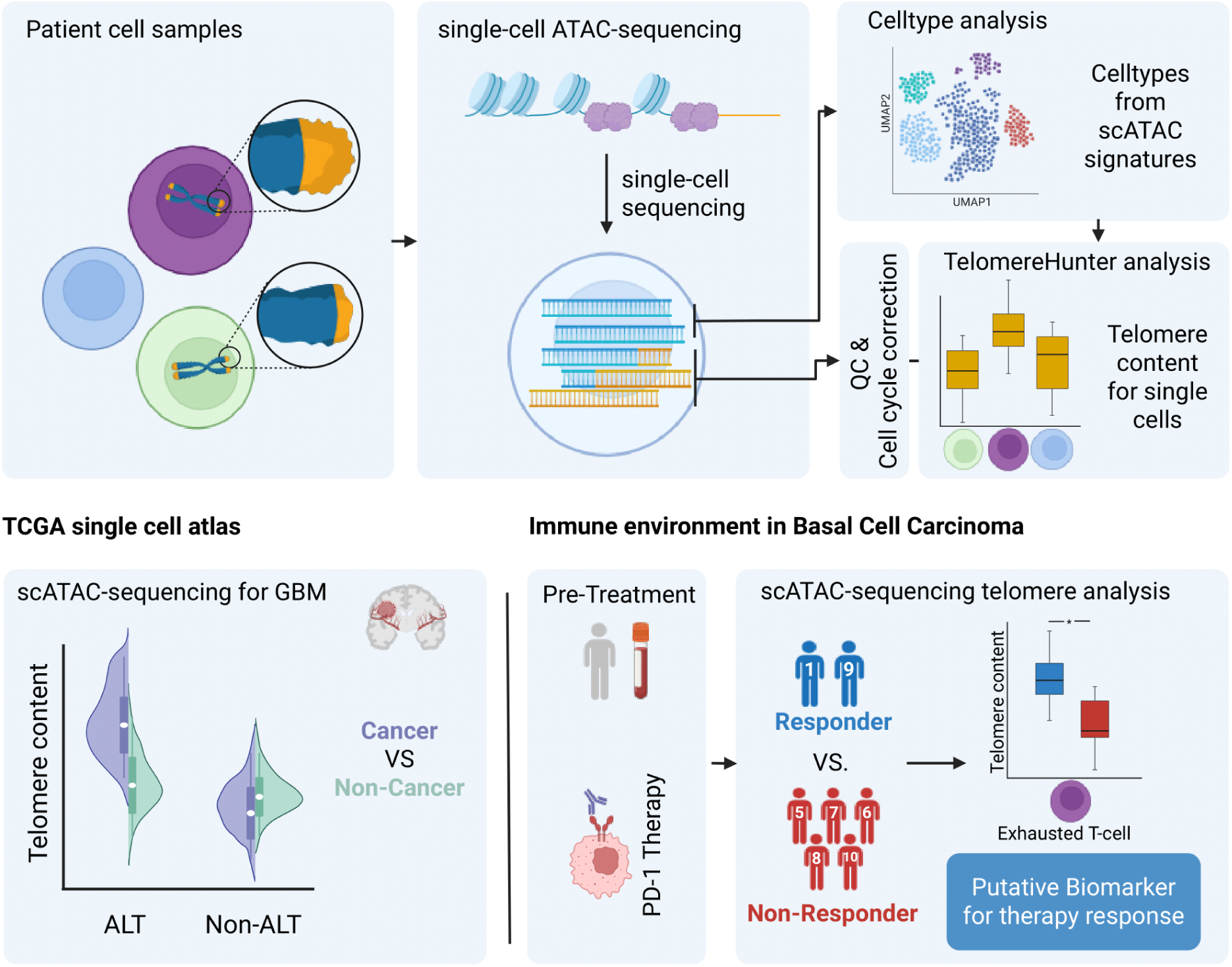
Graphical overview of scATAC-seq–based telomere analysis. Schematic representation of the workflow for estimating telomere length from scATAC-seq data and the downstream analyses conducted using TCGA single-cell atlas and Satpathy et al. datasets.

### Sparseness - Modeling telomeric read detectability in scATAC-seq

As read counts of telomeric sequences are discrete, zeros and low numbers of telomeric reads are only informative on telomere length, if observed in cells with sufficient coverage. If low coverage cells are included, the true telomere length may be underestimated.

To assess how sequencing coverage constrains the detectability of telomeric fragments in single-cell chromatin accessibility profiles, we applied a probabilistic model. The goal was to determine the minimum read depth at which intratelomeric signals can be robustly detected, given that telomeric reads constitute only a minute fraction of the genome and are therefore highly sensitive to both biological variation and technical noise. Conceptually, the model estimates the expected number of telomere-derived reads by integrating assumptions about telomere length, genome size, and technical enrichment. It then quantifies the probability of detecting a defined threshold of intratelomeric reads across different coverage scenarios.

To account for unevenly represented telomeric sequences during scATAC-seq library preparation, we incorporated an amplification bias parameter into the modeling framework. A schematic representation of this effect is provided for reference (Fig. 2a). We modeled the probability of observing at least five intratelomeric reads as a function of sequencing depth (Fig. 2b). At low coverage levels (<10,000 total reads per cell), the probability for individual cells to reach this threshold remains below 50%, even under substantial amplification bias. As sequencing depth increases, detection probabilities rise, exceeding 90% at approximately 30,000 total reads under moderate bias levels (2-3). Beyond this threshold, additional reads result in only marginal improvements in detection.

**Figure 2.**
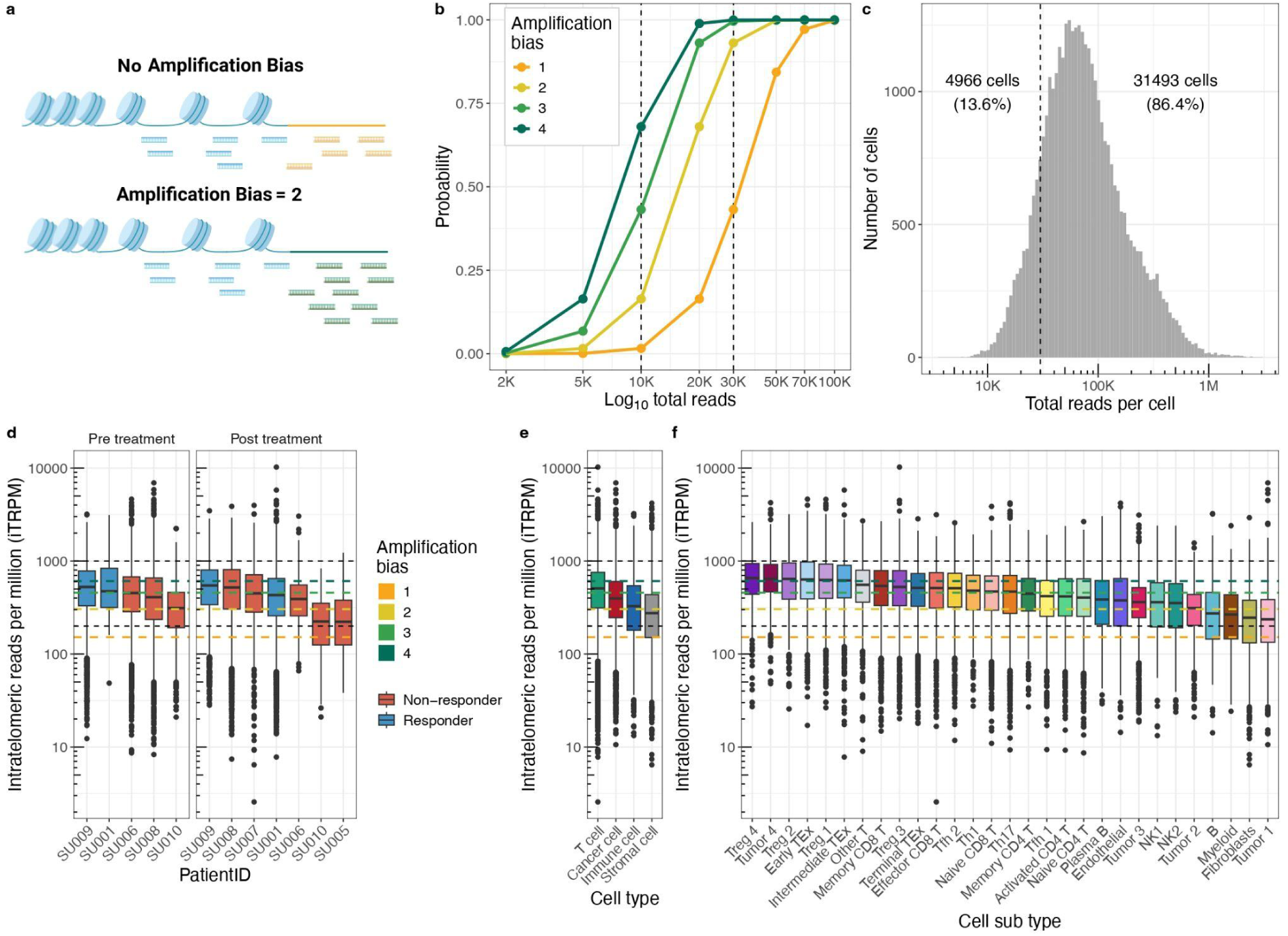
Robustness and sparseness of the ATAC-seq protocol for telomere content estimation. **(a)** Schematic representation of amplification bias in scATAC-seq libraries, where telomeric reads are preferentially enriched, leading to their overrepresentation among total sequencing reads. **(b)** Probability of detecting at least five telomeric reads per cell as a function of sequencing depth (x-axis, log scale) and amplification bias (color scale). Vertical lines mark sequencing depths of 10,000 and 30,000 total reads. Robustness of the ATAC-seq protocol for telomere content estimation. **(c)** Distribution of total reads per single cell in the scATAC-seq dataset. A dashed line indicates the cutoff of 30,000 reads per cell; the number and fraction of cells above and below this threshold are annotated. **(d–f)** Boxplots showing telomeric reads per million total reads for: (d) Patient IDs from Satpathy et al., faceted by pre- and post-treatment and colored by therapy response status (responding vs. non-responding patients), (e) bulk cell types, and (f) cell subtypes. In each plot, dashed black lines indicate iTRPM thresholds of 200 and 1000, and colored dashed lines denote amplification biases from 1 to 4.

To systematically explore parameter space, we assessed detection probabilities under different amplification bias scenarios (Fig. S1). These results demonstrate that telomeric read detection in scATAC-seq is sensitive to sequencing depth and amplification biases. To control for sparseness, we defined a minimum cutoff of 30,000 total reads per cell. This threshold balances the number of informative cells against the total number of included cells. Compared to the recommended minimal coverage for recent single-cell Multiome protocols of 25,000 read-paris per cell, this indicates that usually most, but not all measured cells will be informative. Adopting this criterion reduces noise from poorly covered cells and provides a robust foundation for telomere-focused analyses in scATAC-seq data.

### Representation - Robustness of the ATAC-seq protocol for telomere content estimation

The first analyzed scATAC-seq dataset (Satpathy *et al.*) comprises biopsies obtained from seven patients with BCC, ranging in age from 50 to 75 years^28^. Samples were collected before and at various time intervals after receiving PD-1 immunotherapy. Patient responses to the treatment are available from the original publication (Table S1). Based on the original FACS-sorting fractions (Total fraction: unsorted; T cell fraction: CD45^+^ and CD3^+^; non-T Immune cell fraction: CD45^+^ and CD3^-^; Stromal and Tumor fraction: CD45^-^), and the clustering analysis of the ATAC-seq profiles, we assigned the cells to four groups: Tumor, Immune, Stromal, and T cell. To estimate the telomere content of each cell computationally, we employed TelomereHunter^35^. The derived raw telomere content estimate for each individual cell, represented by their cellular barcode, was filtered by the minimum cutoff of 30,000 total reads per cell, and then corrected for the chromatin state. As a result, 13.6% of cells were excluded from the dataset, leaving 31,493 cells (Fig. 2c).

Next, we tested if the telomerome is underrepresented by this assay. To compute a lower bound for the expected raw telomere content in healthy cells of young individuals, we assume an average telomere length of 10 kbp per chromosome end. Consequently, 960 kbp out of the 6,32 Gbp of genomic DNA in a diploid human genome (average of female genome 6.37 Gbp and male genome 6.27 Gbp) originate from telomeric regions, accounting for approximately 0.0152% of the genome or 152 intra-telomeric reads per million sequenced reads (iTRPM). Recent *in vivo* studies from cell lines suggest a slightly higher interval in human cells between 200 and 1000 iTRPM, respectively 0.02-0.1% of the genomic DNA^39^. Then, we grouped the single-cell telomere content estimates derived from TelomereHunter data according to the available technical and biological annotations, such as their patient IDs (Fig. 2d), bulk cell types (Fig. 2e), and according to the cell type derived from ATAC-seq profiling (Fig. 2f). Across all patient IDs, bulk cell types, and cell subtypes, most distribution medians fall within the 1-4 amplification bias range and between 200-1000 iTRPM. We conclude that telomeres are indeed well represented in ATAC-seq datasets.

### Competing signals - Correcting the telomere signal for chromatin plasticity

Decondensation of chromatin, for instance during the S-phase of the cell cycle, increases the number of potential target sites for the Tn5 transposase that is applied in the ATAC-seq protocol. The resulting competition for the sequencing power acts as a confounder of the *in silico* telomere quantification^34^. To measure this effect, we characterize the global chromatin state by calculating the number of reads per cell, which map to heterochromatic or regulatory genome regions.

In brief, all mapped reads were classified based on the 15 chromatin states of the ChromHMM^40^ reference annotation, and counts were normalized to reads per million sequenced reads (RPM). While reads attributed to regulatory regions displayed a strong positive correlation among each other, they were strongly anti-correlated to the heterochromatin state (Fig. 3a). Next, the seven chromatin states associated with promoters and enhancers were aggregated as regulatory genome regions, and each cell’s global chromatin state was quantified using regulatory and heterochromatin RPM counts, here referred to as the openness score (Fig. 3b).

**Figure 3.**
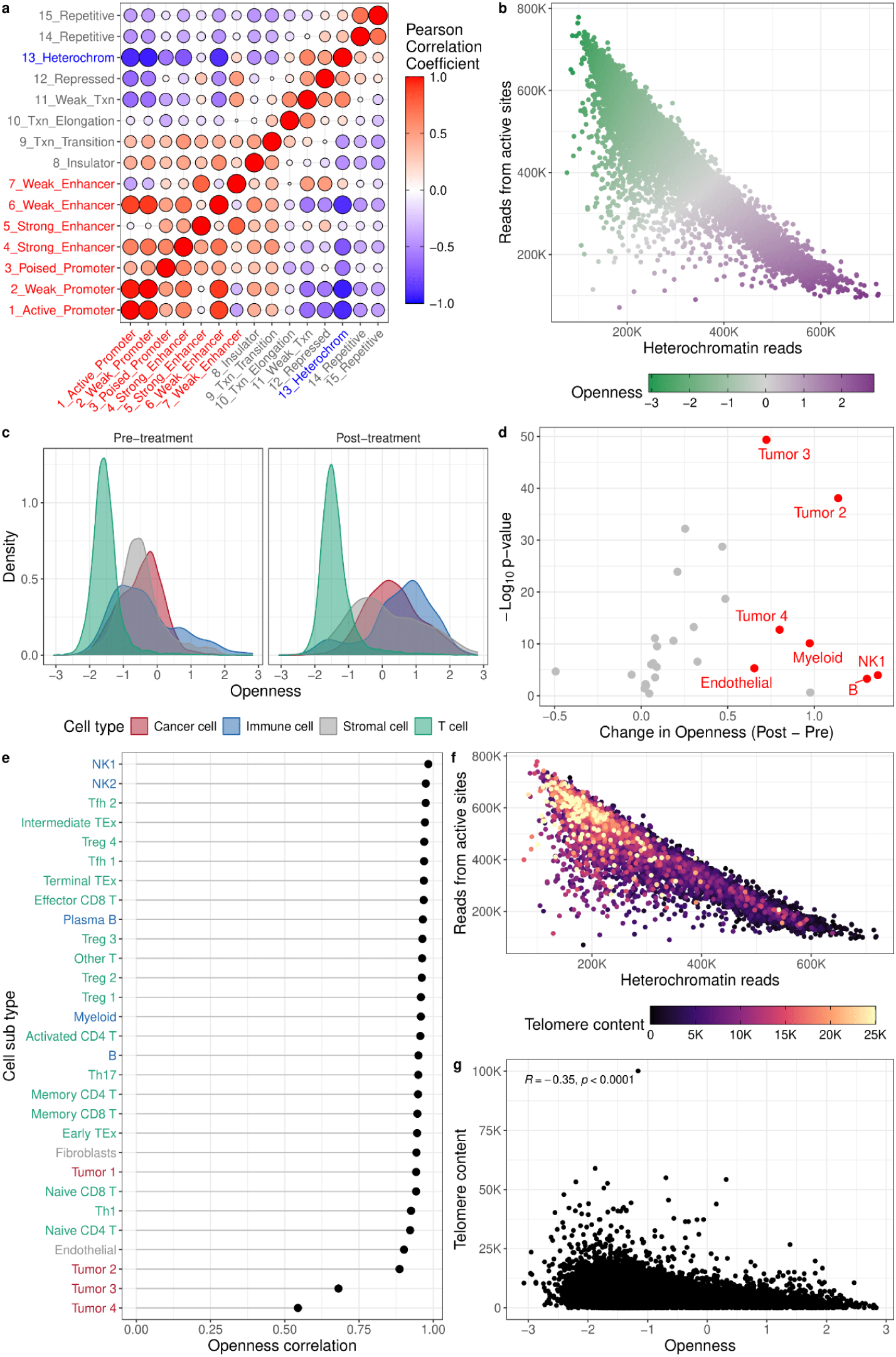
Chromatin condensation state in scATAC-seq from Satpathy et al. **(a)** Pearson correlation matrix of coverage across 15 ChromHMM states in scATAC-seq data. **(b)** Scatter plot of per-cell reads mapping to active regulatory elements (promoters and enhancers, ChromHMM) versus heterochromatin. Cells are colored by the openness score. **(c)** Density distributions of openness scores in cells before and after PD-1 checkpoint blockade, colored by bulk cell type fraction. **(d)** Volcano plot showing differences in openness scores between cell subtype clusters following PD-1 checkpoint blockade. The x axis indicates the change in openness score and the y axis shows statistical significance. Cell subtypes with q < 0.01 are highlighted. **(e)** Openness correlations, displayed as the absolute value of the per-cell correlation between active-site and heterochromatin read coverage (|openness correlation|), summarized by cell subtype and colored by bulk cell type. **(f)** Scatter plot of active-site versus heterochromatin reads per cell, colored by telomere content. **(g)** Relationship between telomere content and openness score at the single-cell level; Spearman correlation coefficient and p-value are indicated.

When examining the distribution of the openness score across the four bulk cell type fractions, T cells exhibited the lowest scores with a very concentrated peak, which remained largely unchanged between pre- and post-treatment. In contrast, Cancer, Immune, and Stromal cell fractions displayed broader distributions, with an overall increase in openness score post-treatment (Fig. 3c). At the resolution of the cell subtype cluster annotation provided by Satpathy *et al.*, this effect was mainly driven by tumor 2, 3, and 4 clusters among cancer cells, endothelial cells among stromal cells, and myeloid, B and NK1 cells among immune cells, all of which showed increased openness post-treatment (Fig. 3d). This suggests that post-treatment, these specific subpopulations may undergo chromatin remodeling, potentially reflecting changes in transcriptional regulation or cellular activation states. Per-cell correlations between active-site and heterochromatin read coverage across cell subtypes were very high, except in tumor clusters 2-4. This suggests that in these tumor cells, the typical relationship between regulatory and heterochromatin regions deviates from that observed in normal cells (Fig. 3e).

We found that cells with lower openness scores exhibited higher telomere content, showing a significant correlation (Fig. 3f,g). After correcting the telomere length estimates for each cell’s openness score, the correlation remained statistically significant, but was reduced to a negligible magnitude (Fig. S2a). Similarly, when binarizing cells into “open” and “closed” chromatin, the previously observed shift toward greater telomere lengths in closed chromatin cells disappeared after applying the openness score correction (Fig. S2b,c).

### TCGA chromatin analysis

We next applied our chromatin openness quantification approach to the scATAC-seq data of the TCGA single-cell atlas. This dataset curated by Sundaram *et al.* (2024)^38^ spans over eight cancer types: colon adenocarcinoma (COAD), breast cancer (BRCA), and lung adenocarcinoma (LUAD), as well as skin cutaneous melanoma (SKCM), kidney renal clear cell carcinoma (KIRC), kidney renal papillary cell carcinoma (KIRP), urothelial bladder carcinoma (BLCA), and glioblastoma multiforme (GBM) and comprises 74 patients. Examining the openness score correlations across different cell types, we observed a pattern broadly similar to the Satpathy *et al.* dataset. While most non-cancer cell types displayed the typical linear relationship seen in the Satpathy data, cancer cells exhibited a distinct, triangular-shaped pattern (Fig. S3). Some other non-cancer cell types, such as B cells, also included a limited number of deviating cells, which may reflect misassigned cell types. We hypothesize that this shift in cancer cells explains why tumor clusters 2-4 in Satpathy *et al.* showed weaker correlations, suggesting the presence of aberrant epigenetic and gene regulatory programs in these cells. Interestingly, when stratifying by cancer type, this pattern was observed to varying degrees in most types, but was largely absent in GBMx cancer cells, indicating that the openness score may capture cancer type-specific alternative epigenetic and gene regulatory programs (Fig. S4). Finally, as with the Satpathy *et al.* data, we observed a negative correlation between openness score and telomere content across most cancer types in TCGA cells, which we corrected for (Fig. S5).

### Identification of ALT-associated telomeric features in single-cell chromatin data

To compare telomere content between cancer and non-cancer cell populations within individual patients, we quantified the adjusted single-cell telomere content for the TCGA scATAC-seq data (Fig. 4a). Most patients displayed reduced telomere content in cancerous cells. This is consistent with literature demonstrating that telomeres in tumor cells are generally shorter than in healthy tissue due to proliferative turnover during tumorigenesis^12^. In contrast, BLCA, BRCA, and GBMx cohorts included samples with significantly higher telomere content in cancer compared to non-cancer cells. Among these, the samples BRCA_14AD76EE_X003, GBMx_A90B18B6_X006, and GBMx_ED12A6C9_X003 (displayed as triangles in Fig 4a) exhibited the largest Cliff’s delta effect sizes with 0.46, 0.82, and 0.71, respectively (Table S2), which motivated their selection for downstream analyses. The distribution of adjusted telomere content within breast cancer and glioblastoma cohorts further confirmed the exceptional telomere content of all three patients. (Fig. 4b). This representation allowed patient-specific inspection of variability within each compartment and revealed a shift toward higher telomere content in the malignant compartment relative to their non-cancer counterparts.

**Figure 4:**
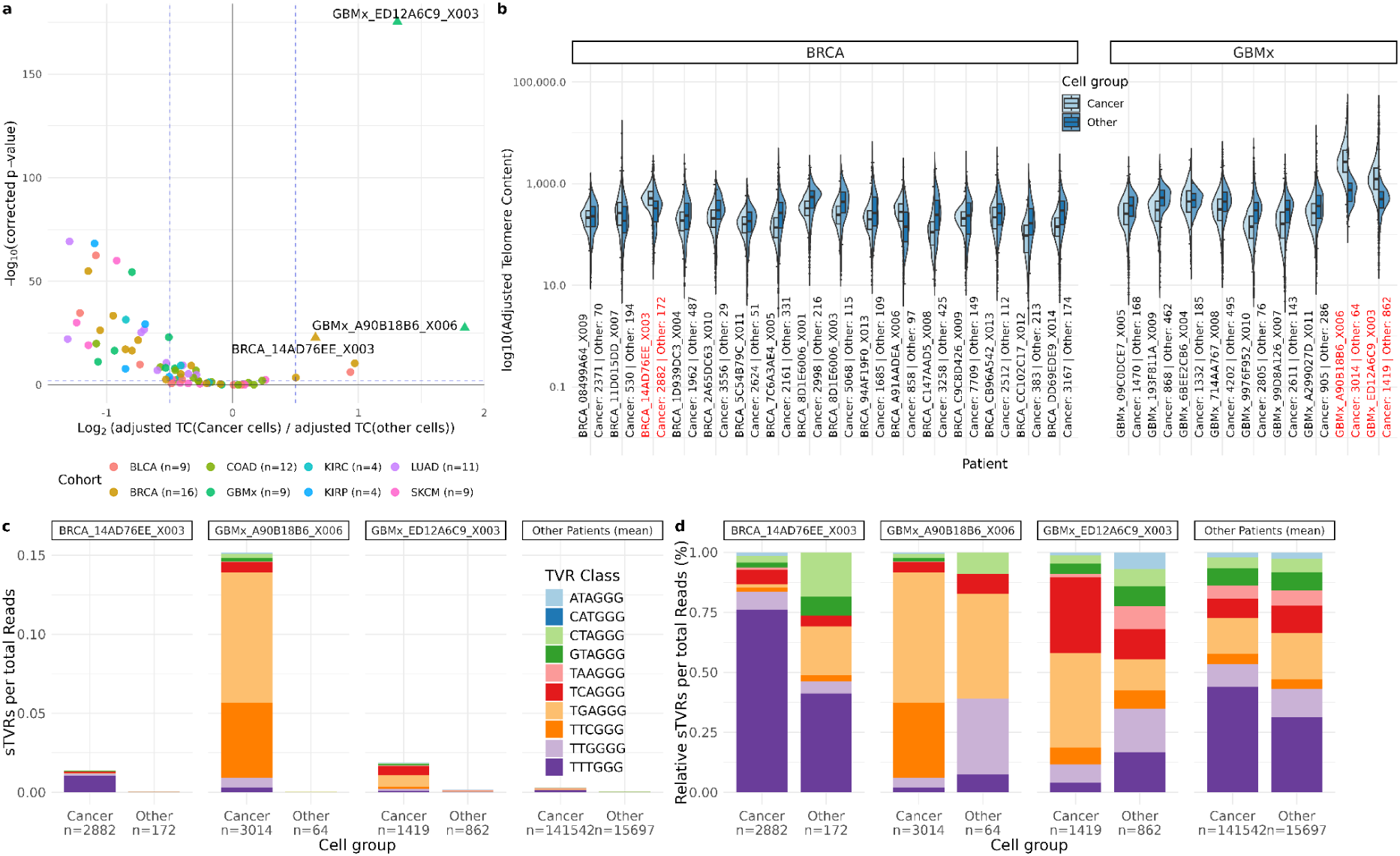
Comparative analysis of telomere content and singleton variant repeats across patient samples and cell types. **(a)** Volcano plot showing differential adjusted telomere content between cancer and non-cancer cell populations (T cell, B cell, Macrophages, Plasma cell, Microglia, Fibroblasts, and Endothelial cells) across individual samples. The x-axis represents the log2 fold change of adjusted telomere content in cancer cells relative to other cell types. P-values are corrected for multiple testing using the Benjamini-Hochberg procedure. Dashed lines indicate the threshold for p values = 0.01 and |log2FC| > 0.5. Triangular points highlight the three patients with the largest effect size (Cliff ’s delta). **(b)** Split violin plots of adjusted telomere content in cancer vs other cells for Breast cancer and Glioblastoma cohorts. Left and right halves represent cancer and non-cancer cells, respectively, with overlaid half-boxplots. Sample labels indicate absolute cell counts for each cell group. Patients of interest BRCA_14AD76EE_X003, GBMx_A90B18B6_X006, and GBMx_ED12A6C9_X003 are highlighted in red. **(c)** Absolute signal of different singleton telomeric variant repeats per total reads in cancer versus other cell populations for the patients of interest and the mean of the remaining patients. **(d)** Relative proportion of singleton telomeric variant repeats per total reads for the same samples and cell groups as in (c). Data are normalized within each cell group to highlight relative TVR composition.

As an additional layer of telomere composition, we profiled singleton telomeric variant repeats (sTVRs) which represent TVRs embedded in canonical telomere sequence^35,22^. They served as informative markers of telomere maintenance mechanisms, as their enrichment patterns can differ between telomerase-driven elongation and alternative lengthening of telomeres (ALT)^19^. The three samples exhibited elevated sTVR signals relative to the mean cancer values of the remaining patients (Fig. 4c). BRCA_14AD76EE_X003 was primarily defined by an isolated enrichment of TTTGGG repeats of more than 10 standard deviations over the cohort mean (z-score > 10) (Table S3, Fig. S8), while other TVR types remained near cohort expectations. This pattern contrasted with the broader and more heterogeneous enrichment observed in the two glioblastoma samples. In one of these, GBMx_A90B18B6_X006, the overall sTVR signal was the highest in the dataset (Fig. 4c), coupled with extensive enrichment across multiple repeat types. TTCGGG, TGAGGG, TTGGGG, TCAGGG, CTAGGG, ATAGGG, and GTAGGG each exceeded a z-score of 10, indicating substantial overrepresentation relative to the baseline distribution. Particularly strong effects wereobserved for TTCGGG (z = 314.56) and TGAGGG (z = 178.01), which surpassed the values measured in other patients. GBMx_ED12A6C9_X003 also showed broad elevation across the same set of sTVRs enriched in GBMx_A90B18B6_X006, with all affected repeats exceeding z-scores of 3. The strongest increases were observed in TCAGGG and TGAGGG with 40.55 and 32.32, respectively, indicating a pronounced but overall less extreme enrichment.

Across all three outlier patients, the enrichment signals increased well above those seen in the remainder of the cohort. These patterns were further emphasized in the relative representation (Fig. 4d), where the three patients of interest stood out in comparison to their own non-cancer compartments, as well as, the averaged malignant and non-malignant values of the other patients.

The distinct TVR patterns observed in the three patients are indicative of differences in telomere maintenance mechanisms. In the breast cancer patient, the prominent enrichment of TTTGGG repeats is consistent with telomerase mediated extension, suggesting active TERT function^24,41,42^. In the two GBMx patients, the enrichment of TGAGGG and TTCGGG repeats aligned with profiles reported for ALT-positive gliomas, supporting the presence of recombination-based telomere elongation in these tumors^43^.

Furthermore, the analysis of subclonal populations in the two Glioblastoma patients GBMx_A90B18B6_X006 and GBMx_193F811A_X009 revealed patient-specific differences in telomere content and sTVR composition. In GBMx_A90B18B6_X006, all three subclones exhibited higher telomere content compared with the other patient (Fig. S9a) and each subclone displayed a similar sTVR profile dominated by TGAGGG and TTCGGG repeats (Fig. S9b). In contrast GBMx_193F811A_X009 showed comparable sTVR signal across its two subclones, which closely resembled the average profiles observed across all patients, indicating a more typical distribution of telomeric variants. These observations highlight that the elevated telomere content and distinct TGAGGG repeat enrichment in GBMx_A90B18B6_X006 are maintained across its subclonal populations.

### Telomere content analysis reveals a distinct PD-1 checkpoint therapy response signature in exhausted T cells

To investigate the interplay between replicative potential and immunotherapy, we returned to the analysis of the Satpathy data. We assessed the potential of telomere content, derived from single-cell profiles, as a biomarker for immunotherapy outcome by resolving measurements by cell type. Prior to treatment, telomere content was significantly higher in cells from responding compared to non-responding patients, both in the bulk T cell fraction(Fig. 5a) and across all cells combined (log_2_FC = 0.12, p < 0.001). After treatment, however, this pattern was reversed, with non-responding cells exhibiting higher telomere content, both in the bulk T cell fraction (Fig. 5a) and across all cells combined (log_2_FC = –0.03, p < 0.01). These observations suggest that pre-treatment telomere content may serve as an indicator of therapeutic success. When analyzing individual cell subtypes in more detail, we found that this effect was primarily driven by exhausted T cells, particularly terminally and intermediately TEx subsets, which showed markedly higher pre-treatment telomere content in responders (Fig. 5b). This indicates that telomere content in TEx cells may be particularly informative for predicting therapy response.

**Figure 5.**
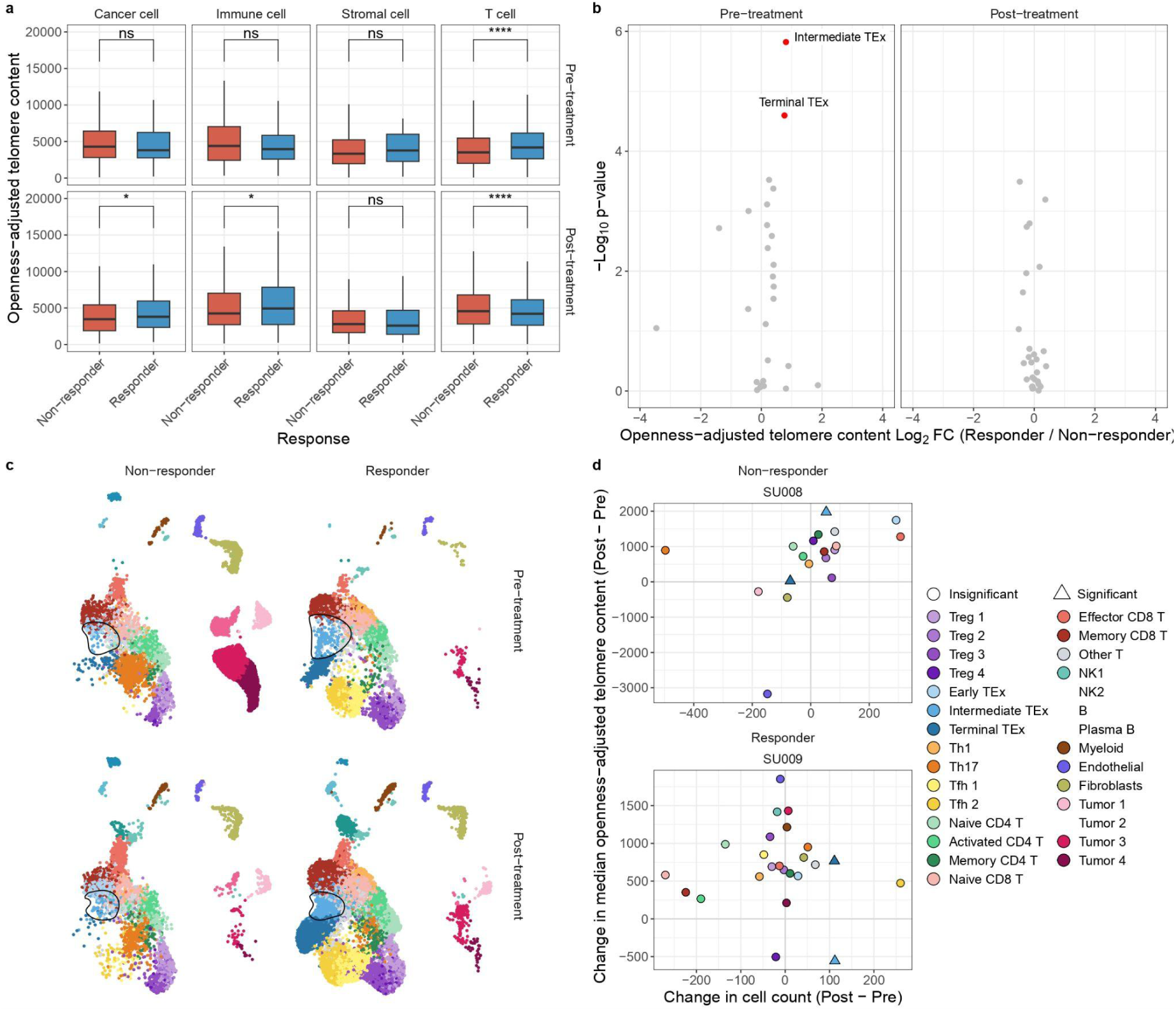
Differences in cell count and telomere content upon PD-1 checkpoint therapy. **(a)** Distribution of single-cell telomere content stratified by therapy response status, treatment time point (pre vs. post), and cell type. Boxplots are colored by therapy response. Significance levels (Wilcoxon test) are shown above each facet. Outliers are not shown. **(b)** Volcano plots of single-cell subtypes comparing telomere content between responding and non-responding patients, using Wilcoxon tests. Separate plots are shown for pre- and post-treatment. Cell subtypes with q-value < 0.01 are annotated. **(c)** UMAP of scATAC-seq cells from Satpathy et al., colored by cell subtype and faceted by therapy response status (responding vs. non-responding) and treatment time point (pre- vs. post-treatment). The location of the Intermediate TEx population is encircled. **(d)** Differences in cell count and median telomere content for each cell subtype between pre- and post-treatment in non-responding patient SU006 and responding patient SU009. Points are colored by cell subtype.

Furthermore, we observed pronounced changes in the abundances of both exhausted T cells and tumor cells during treatment (Fig. 5c). In the responding donor SU009, intermediately exhausted TEx cells expanded their population while simultaneously reducing their telomere content, suggesting proliferative activity (Fig. 5d). Together, these findings demonstrate that telomere content estimates from scATAC-seq can reliably capture cellular replicative potential and support their utility both for quantitative measurements and as biomarkers of immunotherapy response.

## Discussion

In this study, we demonstrate that scATAC-seq can be leveraged for telomere analysis when key technical and biological confounders are explicitly modeled. By addressing read sparsity, telomere representation, and chromatin-state-dependent competition for sequencing signal, we show that telomere content and composition can be quantified at single-cell resolution across cancer and immune cells. This extends telomere analysis to a widely used single-cell modality and enables the integrated interrogation of telomere biology, chromatin state, and cell type.

A fundamental concern for scATAC-seq-based telomere analysis is whether telomeric DNA is adequately represented, given its repetitive sequence and association with shelterin-bound chromatin. Across datasets, patients, and cell types, we observe telomeric read abundances within biologically plausible ranges, consistent with estimates from bulk sequencing and cell line studies^39^. Telomeric reads are not systematically depleted and instead show modest enrichment relative to genomic expectation, indicating that the ATAC-seq protocol captures telomeric DNA sufficiently for comparative analyses. Our probabilistic modeling further shows that sequencing depth is the dominant determinant of telomere signal detectability, allowing us to define a coverage threshold for informative single-cell telomere estimates.

Beyond sparsity, we identify global chromatin condensation as a major and reproducible confounder of telomere content estimation. Cells with globally condensed chromatin, reflected by lower heterochromatin RPM, likely correspond to non-cycling cells (G0) or cycling cells outside of synthesis phase (G1, G2, or M). This interpretation is supported by prior work showing that chromatin accessibility measured by MNase digestion peaks during S phase and decreases during the G phases, with maximal condensation occurring in mitosis^44,45^. Conversely, cells with globally decondensed chromatin and higher heterochromatin RPM are consistent with cells in S phase, when large-scale chromatin decompaction accompanies DNA replication^46^.

Most existing *in silico* approaches that account for cell-cycle effects rely on scRNA-seq-based phase annotation, primarily to control for confounding variation in gene expression^47^. However, such annotations are unavailable for scATAC-seq-only datasets and have been criticized for limited accuracy and robustness, particularly in heterogeneous and cancerous tissues^48^. As discussed by Yakovenko and colleagues, these limitations complicate direct evaluation of the hypothesis that cell-cycle-dependent chromatin condensation biases telomere estimation from scATAC-seq, leading to the conclusion that such measurements are fundamentally unreliable^34^.

By reframing the problem in terms of continuous differences in global chromatin condensation rather than discrete cell-cycle phase labels, our approach circumvents these limitations. Quantifying chromatin state directly from the balance of regulatory and heterochromatic reads allows chromatin-state-dependent bias to be modeled and corrected using scATAC-seq data alone. After correction, the strong correlation between chromatin openness and apparent telomere content is largely reduced, while biologically meaningful differences between cell populations are preserved.

Applying this framework to the TCGA single-cell atlas reveals patient-specific telomere phenotypes consistent with known telomere maintenance mechanisms. While most tumors exhibit reduced telomere content relative to surrounding non-malignant cells, a subset of samples shows elevated telomere content in cancer cells. Analysis of singleton TVRs further distinguishes telomerase-associated patterns from ALT-like profiles. Enrichment of TGAGGG, TCAGGG, TTGGGG and TTCGGG repeats in glioblastoma samples is consistent with prior bulk sequencing studies linking these variants to recombination-based telomere elongation, whereas TTTGGG enrichment in breast cancer aligns with telomerase-driven maintenance^19,22,24^. The preservation of these signatures across subclonal populations suggests that telomere maintenance strategy is a stable and early feature of tumor evolution.

Extending beyond tumor cells, we show that single-cell telomere content captures functionally relevant differences within the immune compartment. Higher pre-treatment telomere content in exhausted T cell subsets, particularly intermediate and terminally exhausted populations, predicts response to PD-1 checkpoint blockade. This is consistent with observations in healthy ageing, where longer telomeres and preserved telomerase inducibility in T cells mark high functional capacity and proliferative fitness^49^. Following treatment, responders exhibit reduced telomere content, consistent with proliferative expansion and telomere attrition in rejuvenated T cells. These observations support telomere content as a quantitative readout of replicative potential within dysfunctional T cell states and highlight its potential utility as a biomarker for immunotherapy responsiveness. As this observation was primarily driven by a single responding patient, caution is required regarding the generalization of this result, but a follow-up on this hypothesis on a larger cohort appears promising.

Several limitations should be considered. scATAC-seq-derived telomere content provides relative rather than absolute estimates and lacks chromosome-end resolution. Detection of telomeric variant repeats is constrained by sequencing depth, limiting per cell analyses. While our chromatin-based correction mitigates cell-cycle-associated bias, definitive validation will require multimodal experimental designs with independent cell-cycle annotation. Nonetheless, the consistency of our findings across datasets, cancer types, and biological contexts demonstrates that telomere analysis from scATAC-seq is feasible and informative when appropriately corrected.

In summary, our work reconciles prior skepticism regarding scATAC-seq-based telomere estimation with empirical evidence and methodological advances. By explicitly modeling chromatin-state effects, we establish scATAC-seq as a practical platform for single-cell telomeromics, enabling integrated studies of telomere maintenance, tumor evolution, and immune function using existing single-cell chromatin datasets.

## Methods

### Data Download and Processing

#### Satpathy et al. (2019) scATAC-seq data

BAM files from the Satpathy *et al.* scATAC-seq dataset^28^ were downloaded from SRA (SRP192525) and converted to fastq files with *bam2fastq* (https://github.com/jts/bam2fastq). Then, “*cellranger-atac count*” from the *Cell Ranger ATAC*^28^ (v. 1.1.0, 10x Genomics) software was used for read filtering, alignment to the reference genome hg19 refdata-cellranger-atac-hg19-1.1.0), peak calling and count matrix generation for each sample. A custom script obtained from 10xGenomics was run for detection of gel bead and barcode multiplets. Barcodes were kept if they passed the following filters: at least 5,000 read-pairs passed read filters; less than 20% read-pairs with low mapping quality “< 0.2 fraction of unmapped + low mapping quality (mapq < 30) + chimeric read-pairs”; less than 90% read pair duplicates; less than 10% read pairs from mitochondrial DNA; no annotation as gel bead or barcode multiplets. Subsequently, sample aggregation was performed using “*cellranger-atac aggr*” without normalization for library size “–normalize = none”. Barcodes with more than 50,000 fragments were subsampled to 50,000 fragments to mitigate the influence of cell count depth on the downstream analysis.

#### TCGA scATAC-seq atlas (2024) data

Single-cell ATAC-seq data was obtained from the Genomics Data Commons portal with approval from the NIH data access committee. As input for processing, we used the pre-assembled fragment files from the repository. We used the standard pipeline from *ArchR*^50^ (v. 1.0.3) under R (v. 4.4.2) on a Debian 12 linux server with 8 threads in parallel. The hg38 genome version was used. Arrow files were generated with the function createArrowFiles with settings of minTSS=4, minFrags=1000, addTileMat=TRUE and addGeneScoreMat=TRUE. *ArchR’s* doublet-removal was performed, and downstream analyses included only cells that had a cell-type annotation in Sundaram *et al.* (2024)^38^ and contained at least 30,000 total reads obtained from running *TelomereHunter*.

### Data visualization and analysis

All data visualizations, data wrangling, and statistical analyses within this project were performed using R (v. 4.4.2, Satpathy dataset and v. 4.4.3, TCGA single-cell atlas). Processing and analysis of scATAC-seq data were carried out using the R packages *ArchR* (v. 1.0.3)^50^, *Seurat* (v. 5.2.1)^51^, and *SingleCellExperiment*^52^. A full list of all R packages used is provided in Supplementary Table 4.

### Statistical Testing

An unpaired Wilcoxon rank-sum test was used to determine the significance of differences between two independent groups. This non-parametric test was chosen because it can handle data that are not normally distributed and because it may be used to compare groups with small sample numbers. In addition to statistical significance testing, effect sizes were quantified using Cliff’s delta to assess the magnitude of differences between groups. P-values obtained from the statistical tests were adjusted using the Benjamini-Hochberg correction to reduce the likelihood of false positives due to multiple testing. A α-level of 0.01 was used as a p- or q-value cutoff. A dedicated description of the statistical procedure used for the sTVR analysis is provided in a separate section below.

### Probabilistic modeling of telomeric read detection in scATAC-seq data

The probability that among the *n* reads of a cell/barcode (sequencing depth) *t* reads are of telomeric origin is modeled a binomial distributed random variable:

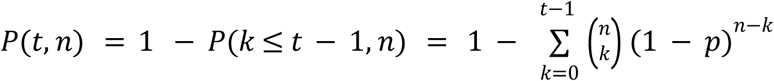

The chance of success p that in an independent draw a read originates from the telomeres is modeled as the ratio of the total telomere length over the genome length times the amplification bias:

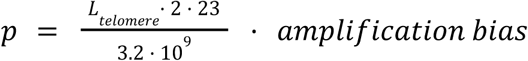

The total telomere length is defined as the product of the average telomere length and the number of chromosome ends. A technical enrichment factor (amplification bias) is included to account for the overrepresentation of telomeric fragments in scATAC-seq libraries. For all simulations, telomere length was fixed at 10,000 bp per chromosomal end, and probabilities were estimated across a grid of sequencing depths and amplification bias values.

### Telomere analysis

The BAM files of the Sapathy dataset were analyzed with *TelomereHunter* v1.1.0. to calculate the raw and GC-corrected telomere content, using the no-control mode^35^. The BAM files of the TCGA dataset were analyzed with *TelomereHunter2* v1.0.0, using the single-cell mode. The parameter --min-reads-per-barcode was set to 30,000. Singleton telomeric variant repeats were quantified at single-cell level also using *TelomereHunter2*. For each cell, the number of sTVRs was normalized by the total number of sequencing reads to account for differences in coverage.

### Enrichment Analysis of sTVRs

The sTVR enrichment analysis of the TCGA dataset was restricted to cancer cells. For each patient, normalized singleton counts were aggregated across all sTVR types and converted to frequencies per 100 cells. These aggregated measures were used to identify patients with unusually high or low overall sTVR abundance. Individuals showing extreme deviations were marked as outliers and omitted from the construction of reference distributions. To assess per-patient shifts in specific sTVR types, the per-100-cell frequencies were standardized across patients using z-scores. For each sTVR type, the mean and standard deviation were calculated only from the non-outlier patients. Two-sided p-values were obtained from the standard normal distribution and corrected for multiple testing using the Benjamini-Hochberg method. sTVR types with |z| > 3 and significant adjusted p-values were considered markedly deviating.

### Openness score and cell-cycle correction

For chromatin state annotation reference in this analysis, the *ChromHMM* annotation for human embryonic stem cell line H1-hESC from ENCODE and Broad Institute was downloaded from the *UCSC Genome Browser*^40^ from the *ChromHMM* segmentation track for the GRCh37/hg19 assembly. This model annotated each 200-base pair segment with one of 15 possible chromatin states^53^. Since the TCGA data was aligned to GRCh38, the *ChromHMM* H1-hESC BED file was converted to GRCh38 via *liftOver*^54^ from GRCh37/hg19.

Then filtered reads were each assigned to a chromatin state to obtain per cell counts for each chromatin state category using bedtools intersect (v. 2.31.1)^55^. To enable comparison between cells, the single-cell read counts were normalized by the total number of reads per cell and multiplied by one million yielding the unit RPM.

To derive a cell-level measure of chromatin condensation and cell cycle state, scATAC-seq reads were aggregated across regulatory regions associated with active chromatin. For each cell *i*, reads mapping to *ChromHMM* states corresponding to active promoter, weak promoter, poised promoter, strong enhancer, and weak enhancer were summed and collectively referred to as active sites (*A _i_*). Reads mapping to the single *ChromHMM* heterochromatin state were summed separately (*H _i_*).

Using these values, we defined a per-cell openness score as

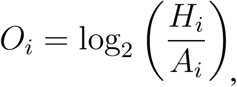

which quantifies the relative enrichment of heterochromatin signal compared to regulatory regions. Higher openness scores indicate increased chromatin accessibility, whereas lower values reflect a more condensed chromatin state.

To correct telomere content for chromatin condensation and cell cycle–associated effects, we modeled the relationship between telomere content and chromatin openness using a linear regression. Let *T_i_* denote the telomere content of cell *i*. We fit the model

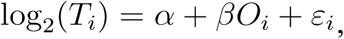

Where *ε_i_* represents the residual term capturing variation in telomere content not explained by chromatin openness.

Corrected telomere content was then defined as

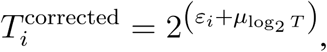

where μ _log₂_ *T* denotes the mean log-transformed telomere content across all cells. This transformation removes systematic variation attributable to chromatin condensation and cell cycle state while preserving the overall scale and distribution of telomere content.

## Code availability

Code and detailed instructions for running *TelomereHunter2* on scATAC-seq data are available at: https://github.com/ferdinand-popp/telomerehunter2. Code used to reproduce the analyses presented in this study is available at: https://github.com/niklas-engel/scATAC_TelomereHunter_2026.

## Author information

## Author contribution

N.L.E. and L.F. conceived the study. L.F. supervised the study. J.G. carried out probabilistic modeling of telomeric read detection. N.L.E. and H.F. performed chromatin condensation analysis. J.G., F.P. and B.B. processed TCGA data, with J.G. performing TCGA telomere analysis with help from N.L.E. L.H., M.S., F.P. and N.L.E. processed the Satpathy dataset, with N.L.E. performing Satpathy telomere analysis. N.L.E., H.F., J.G., and L.F. wrote the manuscript with input from the other authors. All authors read and approved the final manuscript.

## Acknowledgements

This project was funded by the DFG in context of the Forschergruppe “FOR2674: Alters assoziierte epigenetische Veränderungen als therapeutischer Ansatzpunkt in der Behandlung der akuten myeloischen Leukämie”. We thank Cindy Körner for her comments regarding the manuscript. We also thank Daria Morkis for her assistance with data processing. We acknowledge all patients that consented to have their samples analyzed in the Satpathy *et al.* study. The results shown here are in part based upon data generated by The Cancer Genome Atlas (TCGA) Research Network. We further acknowledge the patients and investigators who contributed to the study by Sundaram *et al.* (Science 385, eadk9217, 2024).

## Appendix

**Table S1.**
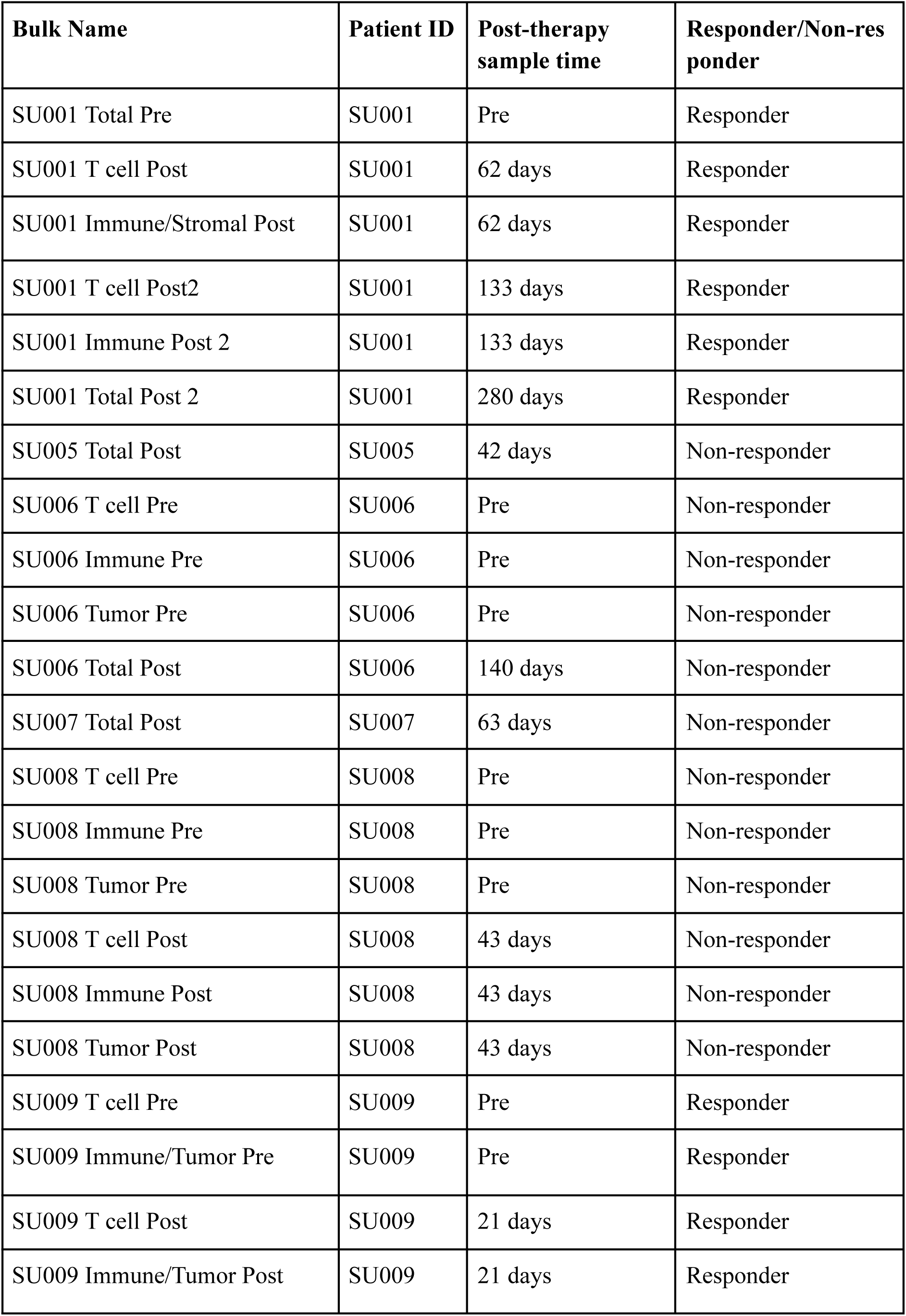
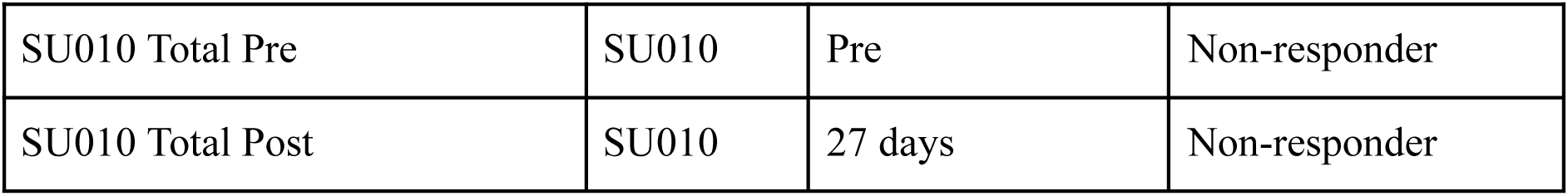
Bulk samples from Satpathy et al. 2019. Overview of the characteristics of each sampled bulk, including the patientID it belongs to, its time of biopsy and the response to PD-1 immunotherapy.

| **Bulk Name** | **Patient ID** | **Post-therapy sample time** | **Responder/Non-responder** |
| --- | --- | --- | --- |
| SU001 Total Pre | SU001 | Pre | Responder |
| SU001 T cell Post | SU001 | 62 days | Responder |
| SU001 Immune/Stromal Post | SU001 | 62 days | Responder |
| SU001 T cell Post2 | SU001 | 133 days | Responder |
| SU001 Immune Post 2 | SU001 | 133 days | Responder |
| SU001 Total Post 2 | SU001 | 280 days | Responder |
| SU005 Total Post | SU005 | 42 days | Non-responder |
| SU006 T cell Pre | SU006 | Pre | Non-responder |
| SU006 Immune Pre | SU006 | Pre | Non-responder |
| SU006 Tumor Pre | SU006 | Pre | Non-responder |
| SU006 Total Post | SU006 | 140 days | Non-responder |
| SU007 Total Post | SU007 | 63 days | Non-responder |
| SU008 T cell Pre | SU008 | Pre | Non-responder |
| SU008 Immune Pre | SU008 | Pre | Non-responder |
| SU008 Tumor Pre | SU008 | Pre | Non-responder |
| SU008 T cell Post | SU008 | 43 days | Non-responder |
| SU008 Immune Post | SU008 | 43 days | Non-responder |
| SU008 Tumor Post | SU008 | 43 days | Non-responder |
| SU009 T cell Pre | SU009 | Pre | Responder |
| SU009 Immune/Tumor Pre | SU009 | Pre | Responder |
| SU009 T cell Post | SU009 | 21 days | Responder |
| SU009 Immune/Tumor Post | SU009 | 21 days | Responder |
| SU010 Total Pre | SU010 | Pre | Non-responder |
| SU010 Total Post | SU010 | 27 days | Non-responder |

**Table S2.** Comparative analysis of openness adjusted telomere content between cancer cells and non-malignant cell population across patients. The columns n_c and n_o denote the number of cancer cells and non-malignant cells (B cells, T cells, Macrophages, Microglia, Plasma cells, Fibroblasts, Endothelial cells), respectively. TC_c and TC_o represent the median openness adjusted telomere content in cancer and non-malignant cells; these values were used to compute the log2-fold change (log2_fc). Effect sizes (eff.size) were calculated using Cliff ’s delta, and p-values were obtained from a two-sided Wilcoxon rank-sum test and adjusted for multiple testing using the Benjamini-Hochberg method. Values are sorted by descending effect sizes.

| Sample | n_c | n_o | TC_c | TC_o | eff.size | log2_fc | p_value_adj |
| --- | --- | --- | --- | --- | --- | --- | --- |
| GBMx_A90B18B6_X006 | 3014 | 64 | 2666 | 742.2 | 0.8154 | 1.845 | 2.929e-28 |
| GBMx_ED12A6C9_X003 | 1419 | 862 | 1226 | 494.4 | 0.709 | 1.31 | 5.869e-176 |
| BRCA_14AD76EE_X003 | 2882 | 172 | 515.1 | 326.2 | 0.4594 | 0.659 | 1.56e-23 |
| BRCA_A91AADEA_X006 | 858 | 97 | 278.2 | 141.9 | 0.418 | 0.9708 | 3.726e-11 |
| BLCA_1067A79F_X003 | 4697 | 93 | 323.1 | 168.8 | 0.3075 | 0.9371 | 6.698e-07 |
| BRCA_11D015DD_X007 | 530 | 194 | 263.7 | 186.1 | 0.1815 | 0.5026 | 0.0002888 |
| SKCM_211D9CF4_X004 | 259 | 223 | 197.1 | 164.4 | 0.1585 | 0.2619 | 0.003849 |
| COAD_1E46B832_X004 | 1181 | 145 | 352.4 | 299.4 | 0.1408 | 0.2353 | 0.007721 |
| COAD_32AA8B86_X013 | 1205 | 172 | 291.8 | 267.5 | 0.08521 | 0.1255 | 0.087 |
| BLCA_CC57A8ED_X006 | 2649 | 153 | 261.4 | 245.9 | 0.04834 | 0.08781 | 0.3422 |
| COAD_1B956E80_X005 | 188 | 30 | 207.2 | 184.3 | 0.03333 | 0.169 | 0.8038 |
| SKCM_8F708E04_X011 | 594 | 353 | 224.5 | 210 | 0.01251 | 0.09688 | 0.7906 |
| BLCA_761A45CF_X011 | 3876 | 186 | 241.2 | 230 | 0.001709 | 0.06865 | 0.9686 |
| BLCA_C1820F13_X010 | 2061 | 110 | 280.3 | 278.8 | -0.00449 | 0.007477 | 0.9497 |
| COAD_74EF44ED_X013 | 935 | 126 | 312.1 | 302.6 | -0.008183 | 0.04448 | 0.9063 |
| GBMx_6BEE2CB6_X004 | 1332 | 185 | 446.6 | 474.2 | -0.01866 | -0.08663 | 0.7306 |
| SKCM_F05B8E69_X007 | 1752 | 175 | 322.8 | 386.6 | -0.05066 | -0.2602 | 0.3016 |
| BRCA_08499A64_X009 | 2371 | 70 | 213.5 | 228.2 | -0.07293 | -0.0965 | 0.3293 |
| COAD_0914606C_X006 | 2386 | 310 | 211 | 236.8 | -0.07422 | -0.1662 | 0.04338 |
| LUAD_E25DF7DA_X010 | 109 | 506 | 314.9 | 359.7 | -0.09066 | -0.1918 | 0.1618 |
| BRCA_C9C8D426_X009 | 7709 | 149 | 208.3 | 236.9 | -0.09314 | -0.186 | 0.06436 |
| KIRC_F62461A5_X012 | 1263 | 117 | 223.9 | 255.2 | -0.1102 | -0.1888 | 0.06201 |
| SKCM_0852FA43_X005 | 1048 | 47 | 154 | 196.6 | -0.1199 | -0.353 | 0.1899 |
| BRCA_5C54B79C_X011 | 2624 | 51 | 159.5 | 184.1 | -0.1277 | -0.2069 | 0.1411 |
| BRCA_CB96A542_X013 | 2512 | 112 | 217.4 | 262.4 | -0.1446 | -0.2716 | 0.01262 |
| COAD_B5AE6F25_X011 | 1641 | 141 | 209.1 | 285.4 | -0.1481 | -0.4488 | 0.00489 |
| SKCM_F15664E6_X012 | 366 | 46 | 224.7 | 299.8 | -0.1535 | -0.4163 | 0.1092 |
| COAD_E51C5352_X009 | 1884 | 475 | 288.2 | 352.4 | -0.1608 | -0.2905 | 1.097e-07 |
| COAD_0B7D5055_X014 | 2034 | 77 | 242.4 | 292.7 | -0.1774 | -0.2721 | 0.01103 |
| BRCA_1D939DC3_X004 | 1962 | 487 | 187.5 | 235.5 | -0.1854 | -0.3283 | 5.152e-10 |
| LUAD_CA6F245B_X006 | 370 | 398 | 231.4 | 282.5 | -0.1863 | -0.2874 | 1.353e-05 |
| COAD_BA3E7A65_X007 | 1647 | 115 | 223.5 | 277.4 | -0.2064 | -0.3113 | 0.0003289 |
| SKCM_F318F3E5_X009 | 3339 | 70 | 163.1 | 227.5 | -0.214 | -0.4803 | 0.003141 |
| LUAD_ECBD63B4_X004 | 720 | 492 | 331.4 | 435.9 | -0.2164 | -0.3954 | 3.646e-10 |
| LUAD_18545E4A_X005 | 7782 | 346 | 195 | 281 | -0.2169 | -0.5272 | 2.181e-11 |
| KIRC_0BCD21B9_X007 | 960 | 113 | 112.9 | 150.6 | -0.2237 | -0.4165 | 0.0001595 |
| GBMx_A299027D_X011 | 905 | 286 | 263.1 | 361.6 | -0.2288 | -0.4586 | 1.088e-08 |
| LUAD_202446E0_X002 | 221 | 377 | 188.1 | 239.9 | -0.2293 | -0.3511 | 5e-06 |
| BLCA_6728BCB4_X012 | 2747 | 58 | 175.5 | 231.5 | -0.2368 | -0.3998 | 0.002977 |
| KIRP_5C0BAEF0_X010 | 518 | 96 | 160.6 | 227.7 | -0.2549 | -0.5032 | 0.0001193 |
| COAD_57255B8E_X010 | 1685 | 194 | 164.7 | 245.2 | -0.2659 | -0.5739 | 2.796e-09 |
| BRCA_94AF19F0_X013 | 1685 | 109 | 198.4 | 264 | -0.2661 | -0.4117 | 5.427e-06 |
| GBMx_714AA767_X008 | 4202 | 495 | 312.7 | 444.2 | -0.2794 | -0.5067 | 1.02e-23 |
| GBMx_09C0DCE7_X005 | 1470 | 168 | 255.9 | 370 | -0.2794 | -0.5319 | 6.074e-09 |
| BLCA_3E71B261_X005 | 483 | 7 | 213.3 | 298.7 | -0.2967 | -0.4855 | 0.203 |
| COAD_D0F82B03_X003 | 5013 | 108 | 246.5 | 337.9 | -0.3058 | -0.4549 | 9.884e-08 |
| KIRC_7D6A394E_X005 | 1335 | 160 | 281.5 | 374.5 | -0.3079 | -0.4117 | 4.355e-10 |
| BRCA_2A65DC63_X010 | 3556 | 29 | 207.2 | 299.2 | -0.3366 | -0.5301 | 0.00269 |
| LUAD_84B38240_X008 | 614 | 836 | 176.2 | 286.8 | -0.3375 | -0.7026 | 2.106e-27 |
| LUAD_202446E0_X007 | 768 | 577 | 137.7 | 227.8 | -0.3401 | -0.7264 | 5.317e-26 |
| LUAD_A1ABC0B2_X009 | 253 | 127 | 178.4 | 270 | -0.3432 | -0.5979 | 9.487e-08 |
| BRCA_8D1E6006_X001 | 2998 | 216 | 326.7 | 549.8 | -0.4005 | -0.7511 | 2.616e-22 |
| BLCA_CC5A9968_X004 | 1188 | 95 | 173 | 288.2 | -0.4031 | -0.7365 | 1.491e-10 |
| SKCM_398F831B_X014 | 286 | 405 | 102.8 | 227.5 | -0.413 | -1.146 | 7.13e-20 |
| BRCA_7C6A3AE4_X005 | 2161 | 331 | 136.9 | 264.3 | -0.4209 | -0.949 | 4.092e-34 |
| BRCA_CC102C17_X012 | 383 | 213 | 95.19 | 165.7 | -0.4237 | -0.7993 | 2.9e-17 |
| GBMx_99D8A126_X007 | 2611 | 143 | 162.6 | 311.7 | -0.425 | -0.9392 | 3.02e-17 |
| KIRP_DFEC4B50_X009 | 2175 | 276 | 229.5 | 371.4 | -0.4262 | -0.6947 | 4.537e-30 |
| KIRP_3AB3B701_X011 | 2274 | 60 | 164.5 | 296.9 | -0.4352 | -0.8516 | 1.681e-08 |
| KIRC_5786B3A8_X006 | 4537 | 252 | 201.4 | 362.5 | -0.4477 | -0.8477 | 3.256e-32 |
| SKCM_FE986D7E_X006 | 2483 | 526 | 182.7 | 346 | -0.4601 | -0.9211 | 1.007e-60 |
| GBMx_9976F952_X010 | 2805 | 76 | 142.1 | 298.4 | -0.4703 | -1.07 | 6.904e-12 |
| BRCA_C147AAD5_X008 | 3258 | 425 | 110.3 | 244.3 | -0.4726 | -1.147 | 1.226e-55 |
| BRCA_8D1E6006_X003 | 5068 | 115 | 243.9 | 440.2 | -0.4766 | -0.852 | 6.623e-18 |
| BRCA_DD69EDE9_X014 | 3167 | 174 | 142.8 | 296.2 | -0.4912 | -1.053 | 4.289e-27 |
| COAD_C4C5A550_X010 | 2646 | 119 | 185.3 | 392.3 | -0.5107 | -1.083 | 1.325e-20 |
| GBMx_193F811A_X009 | 868 | 462 | 302.4 | 526.4 | -0.5253 | -0.7994 | 3.708e-55 |
| BLCA_795D3511_X008 | 2519 | 367 | 126.4 | 268.6 | -0.547 | -1.087 | 3.272e-63 |
| LUAD_8B7E9FCA_X014 | 969 | 107 | 95.11 | 236 | -0.5868 | -1.311 | 7.781e-23 |
| LUAD_9B9C5C6D_X003 | 1772 | 355 | 113.4 | 278.6 | -0.6004 | -1.296 | 6.193e-70 |
| KIRP_E65C8B50_X008 | 3706 | 301 | 143.9 | 307.9 | -0.6137 | -1.097 | 5.651e-69 |
| BLCA_6E1FF9EA_X007 | 2131 | 148 | 118.8 | 275.4 | -0.6185 | -1.213 | 1.935e-35 |
| SKCM_FDA487D2_X008 | 1238 | 111 | 153 | 361.3 | -0.6692 | -1.24 | 9.151e-31 |
| LUAD_3AD9F8C2_X011 | 0 | 429 | NA | NA | NA | NA | NA |

**Table S3.** Patient-Level Summary of Singleton Z-Scores. Z-Transformed sTVR-Values per patient, calculated using only cancer cells and excluding the three outliers samples from mean and standard deviation estimation. The table includes patients with |z-score| > 3 and corresponding two-tailed p-values which were adjusted for multiple testing using the Benjamini-Hochberg procedure (p_adj). Values are sorted by cancer type and descending z-score per patient.

| Sample | TVR_type | sum_per100cells | zscore | p_adj |
| --- | --- | --- | --- | --- |
| BLCA_C1820F13_X010 | TCAGGG | 6e-05 | 4.74252 | 8.6672e-05 |
| BLCA_C1820F13_X010 | ATAGGG | 1e-05 | 3.87654 | 3.6637e-03 |
| BLCA_CC5A9968_X004 | TAAGGG | 3e-05 | 3.30726 | 2.4759e-02 |
| BRCA_14AD76EE_X003 | TTTGGG | 0.00037 | 10.00497 | < 6.3061e-14 |
| BRCA_C9C8D426_X009 | TTCGGG | 2e-05 | 3.32316 | 2.4364e-02 |
| GBMx_A90B18B6_X006 | TTCGGG | 0.00156 | 314.55571 | < 6.3061e-14 |
| GBMx_A90B18B6_X006 | TGAGGG | 0.00274 | 178.01143 | < 6.3061e-14 |
| GBMx_A90B18B6_X006 | TTGGGG | 0.00021 | 22.49635 | < 6.3061e-14 |
| GBMx_A90B18B6_X006 | TCAGGG | 0.00021 | 19.79771 | < 6.3061e-14 |
| GBMx_A90B18B6_X006 | ATAGGG | 4e-05 | 11.22756 | < 6.3061e-14 |
| GBMx_A90B18B6_X006 | CTAGGG | 8e-05 | 9.90331 | < 6.3061e-14 |
| GBMx_A90B18B6_X006 | GTAGGG | 7e-05 | 8.9722 | < 6.3061e-14 |
| GBMx_ED12A6C9_X003 | TCAGGG | 0.00042 | 40.5513 | < 6.3061e-14 |
| GBMx_ED12A6C9_X003 | TGAGGG | 0.00052 | 32.32238 | < 6.3061e-14 |
| GBMx_ED12A6C9_X003 | TTCGGG | 9e-05 | 17.87129 | < 6.3061e-14 |
| GBMx_ED12A6C9_X003 | TTGGGG | 1e-04 | 9.90263 | < 6.3061e-14 |
| GBMx_ED12A6C9_X003 | GTAGGG | 6e-05 | 7.50168 | 3.1870e-12 |
| GBMx_ED12A6C9_X003 | CTAGGG | 5e-05 | 4.95257 | 3.4371e-05 |
| GBMx_ED12A6C9_X003 | ATAGGG | 2e-05 | 4.27251 | 7.4698e-04 |
| KIRC_F62461A5_X012 | TTCGGG | 2e-05 | 3.04889 | 5.8040e-02 |
| KIRP_DFEC4B50_X009 | TAAGGG | 3e-05 | 3.51096 | 1.3334e-02 |
| LUAD_202446E0_X002 | GTAGGG | 3e-05 | 3.41603 | 1.8151e-02 |
| SKCM_211D9CF4_X004 | CTAGGG | 5e-05 | 4.88399 | 4.5534e-05 |
| SKCM_8F708E04_X011 | TTGGGG | 5e-05 | 4.18005 | 1.0638e-03 |
| SKCM_FE986D7E_X006 | TTCGGG | 2e-05 | 3.82202 | 4.3481e-03 |
| SKCM_FE986D7E_X006 | CTAGGG | 4e-05 | 3.63641 | 8.6493e-03 |

**Figure S1.**
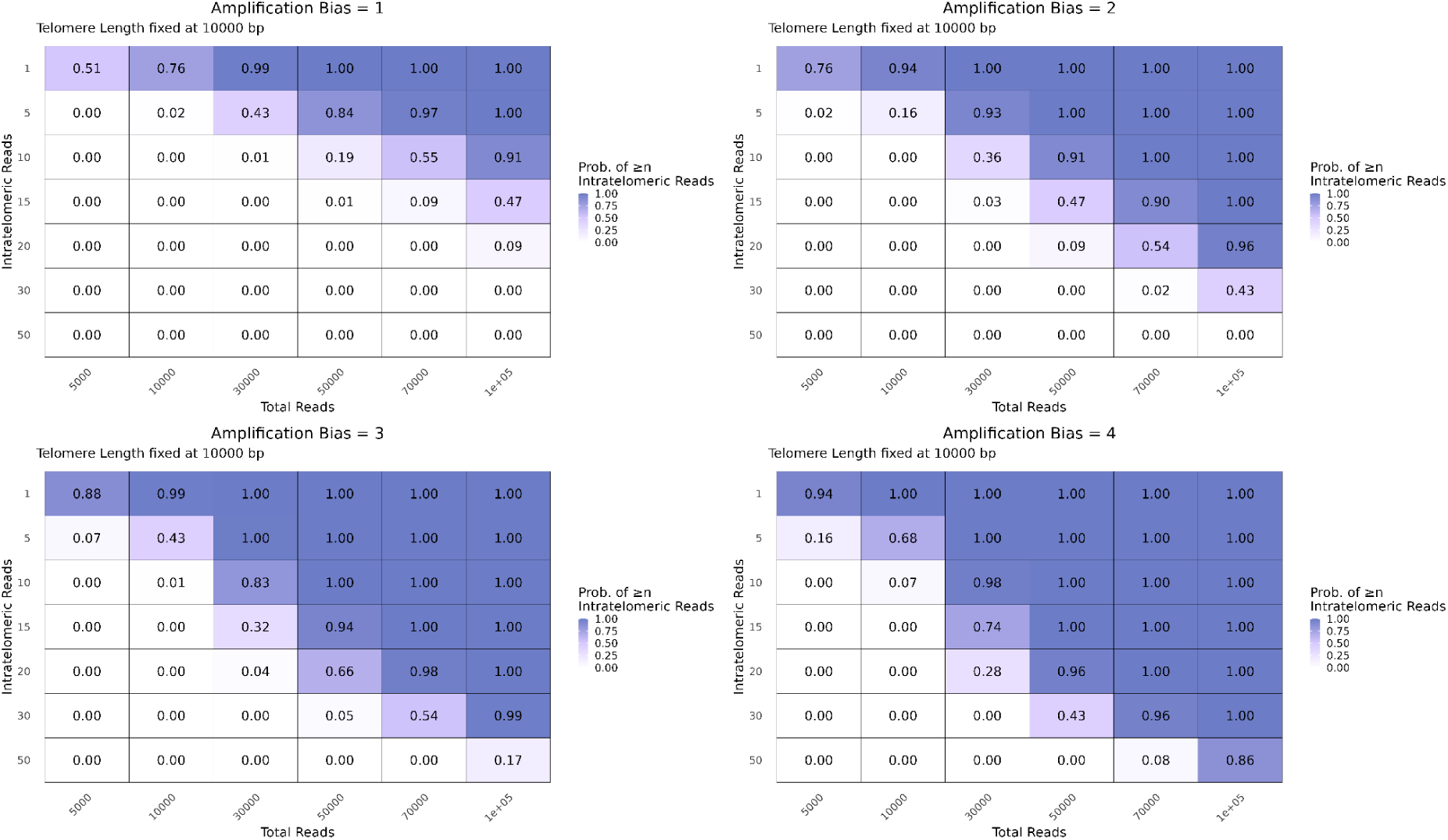
Probability of detecting intratelometric reads across sequencing depth and amplification bias. Heatmaps showing the probability of observing at least n telomeric reads for a fixed telomere length of 10,000 bp under four amplification bias conditions (bias = 1-4). Rows indicate the expected number of intratelomeric reads, while columns represent the total number of sequencing reads. Color intensity reflects the probability values, with darker shades corresponding to higher probabilities.

**Figure S2.**
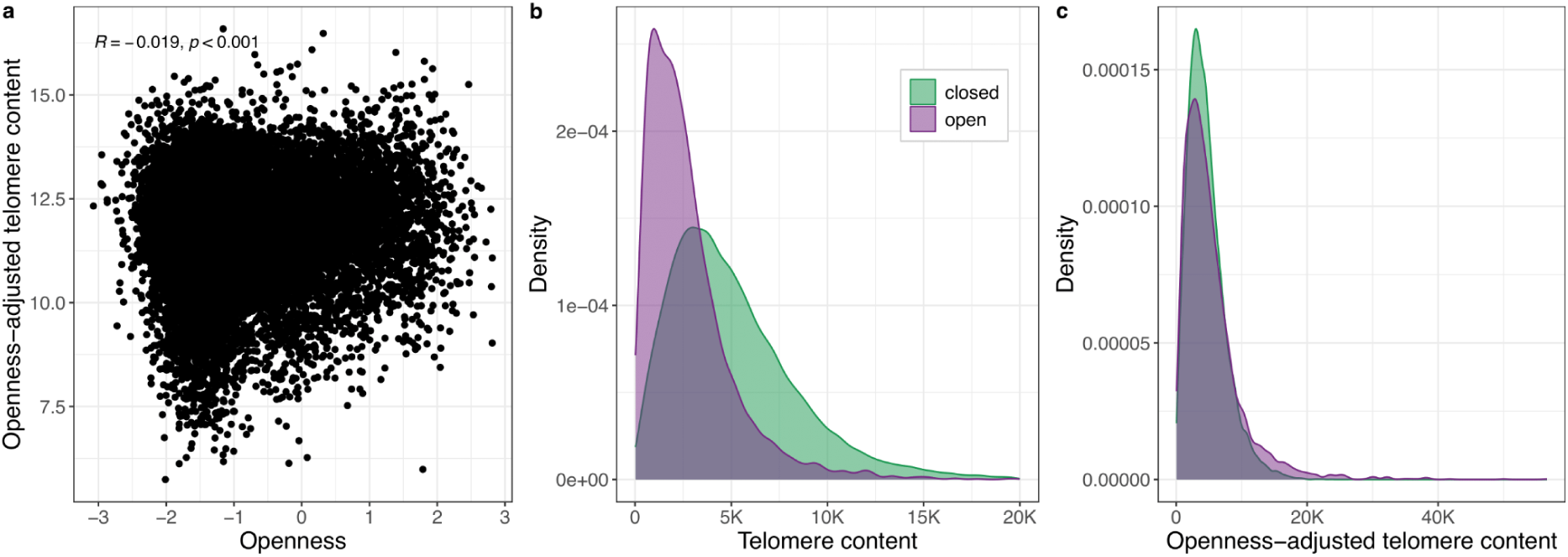
Correcting telomere content for chromatin condensation. (a) Relationship between telomere content adjusted for chromatin openness and the openness score at the single-cell level; Spearman correlation coefficient and P value are indicated. (b,c) Density distributions of (b) telomere content and (c) openness-adjusted telomere content. Cells are stratified by chromatin state, classified as open (openness score > 0) or closed (openness score < 0).

**Figure S3.**
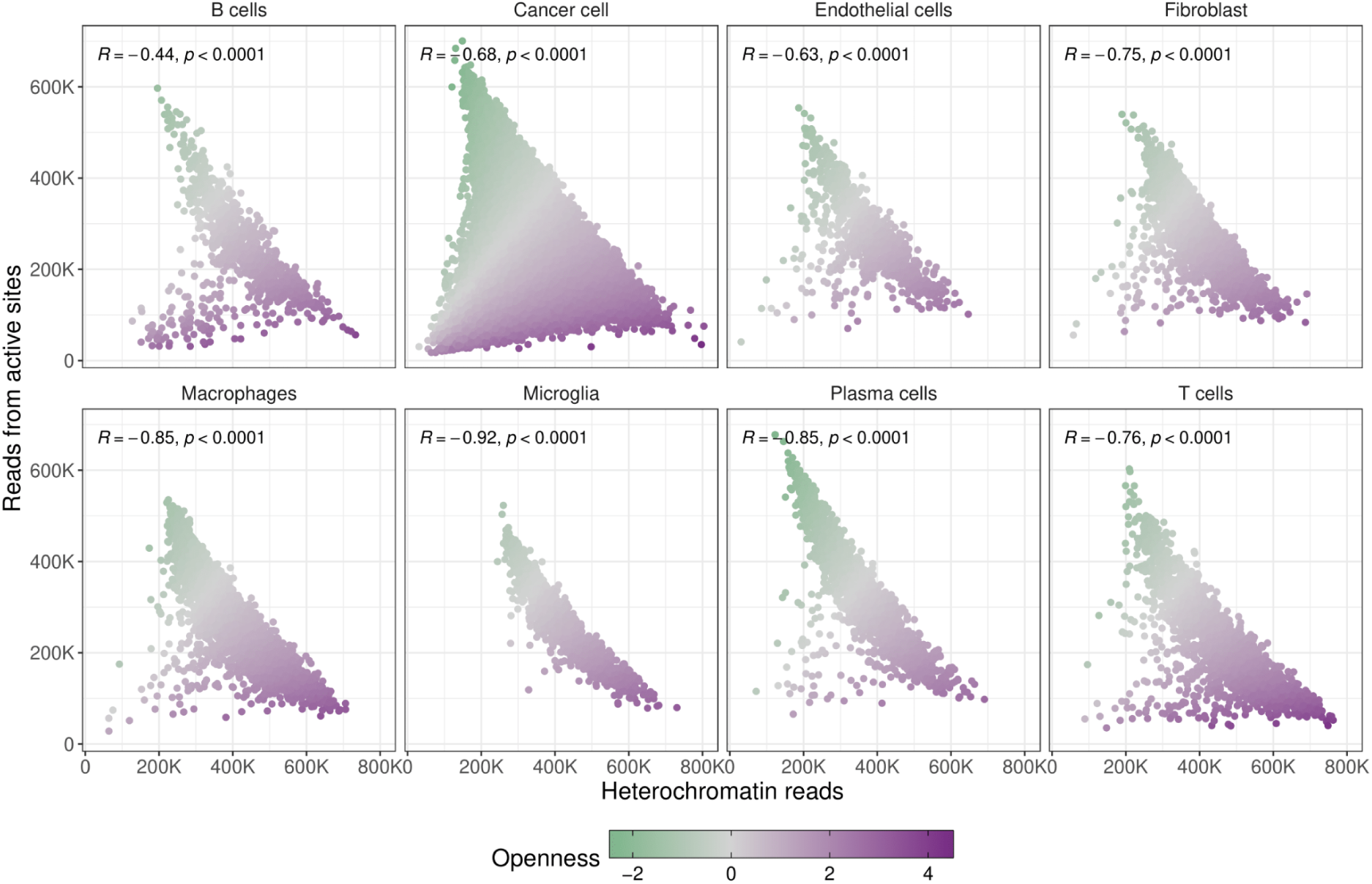
Chromatin condensation across TCGA cell types. Scatter plot of per-cell reads mapping to active regulatory elements (promoters and enhancers, ChromHMM) versus heterochromatin in scATAC-seq data. Cells are colored by the openness score, with each facet representing a cell type from the TCGA scATAC-seq atlas.

**Figure S4.**
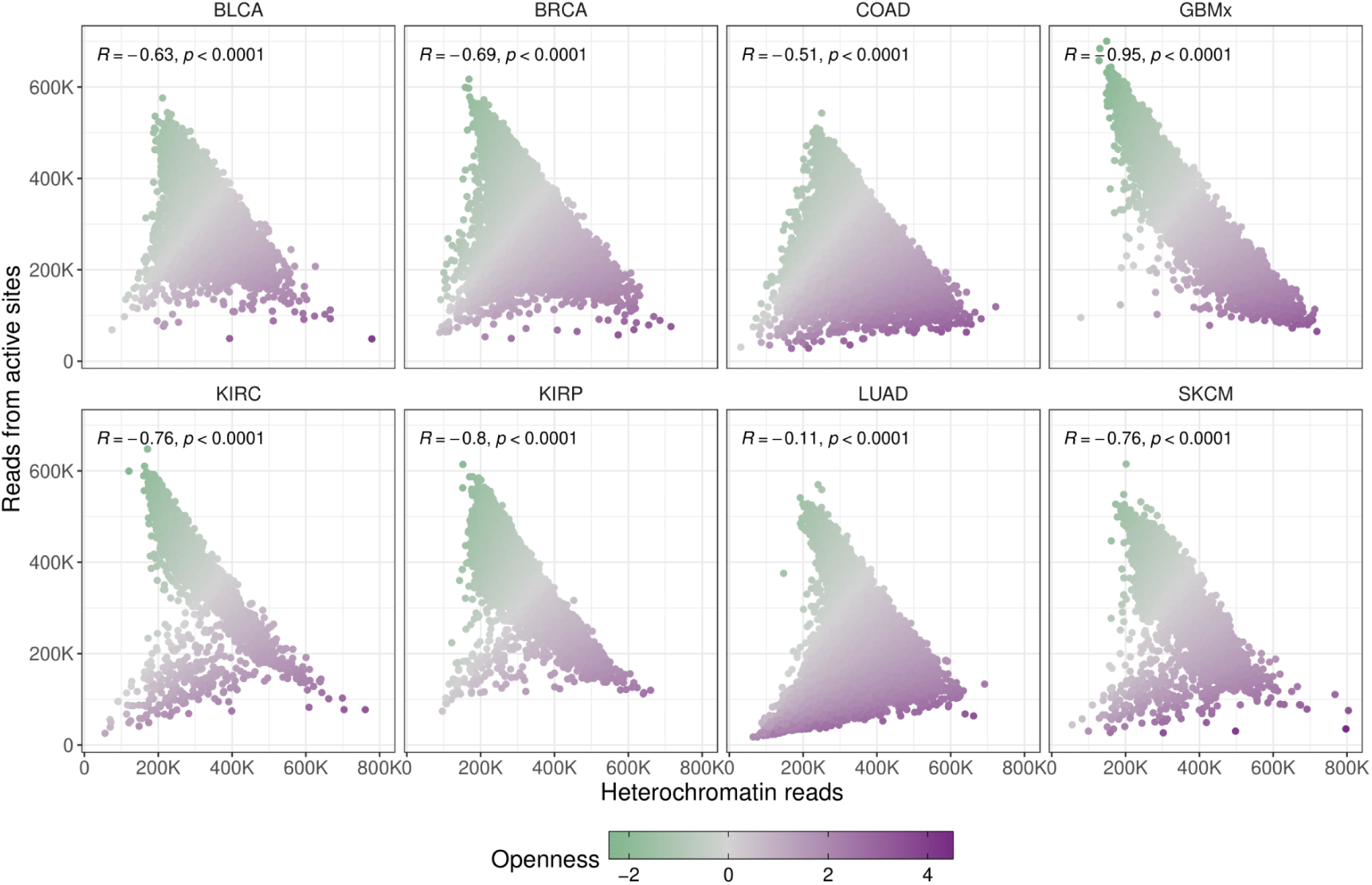
Chromatin condensation across TCGA cancer types. Scatter plot of per-cell reads mapping to active regulatory elements (promoters and enhancers, ChromHMM) versus heterochromatin in scATAC-seq data. Cells are colored by the openness score, with each facet representing a cancer type from the TCGA scATAC-seq atlas.

**Figure S5.**
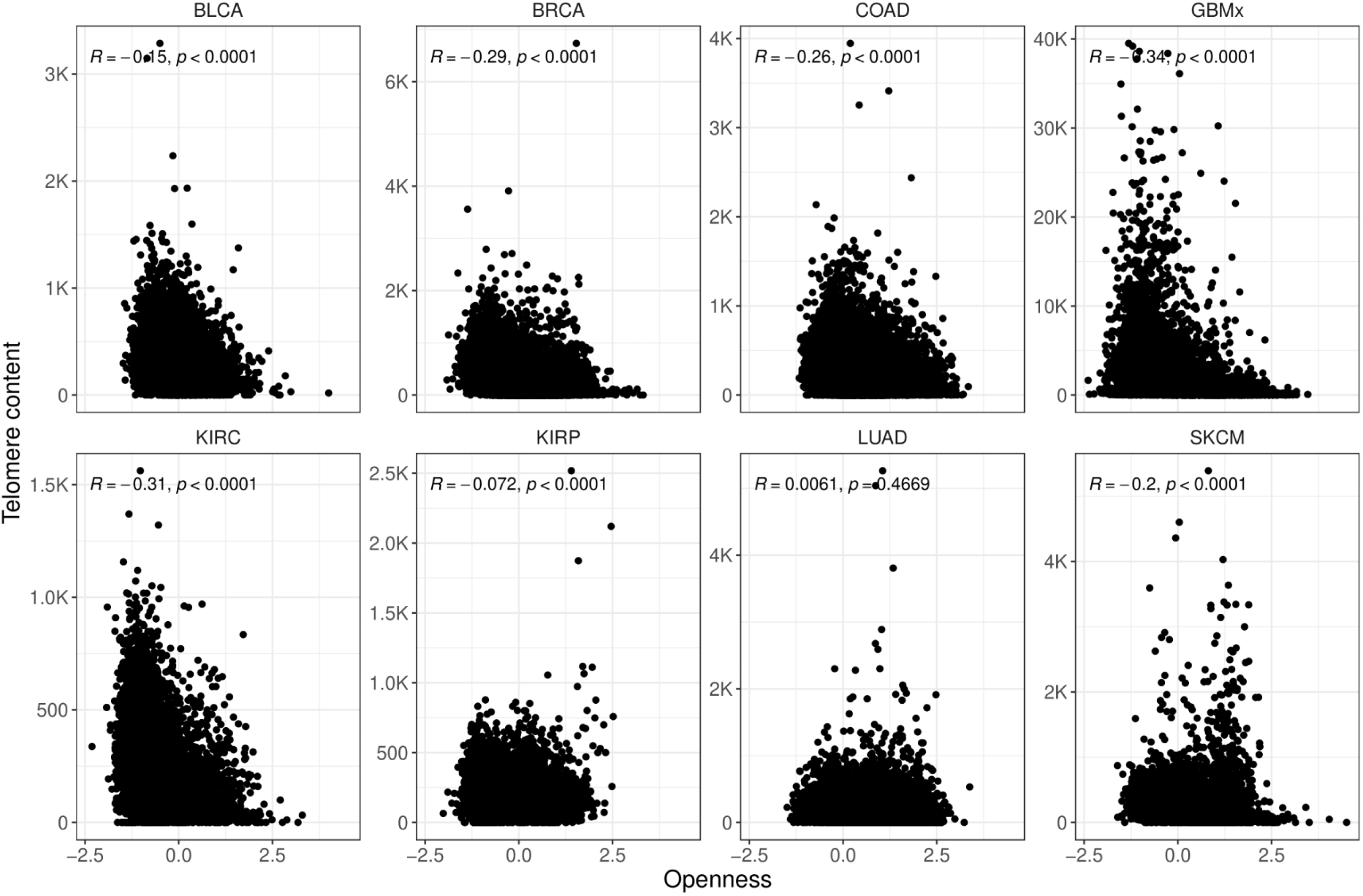
Correcting telomere content in TCGA scATAC-seq data. Relationship between telomere content and openness score at the single-cell level; Spearman correlation coefficient and p-value are indicated. Each facet represents a cancer type from the TCGA scATAC-seq atlas.

**Figure S6.**
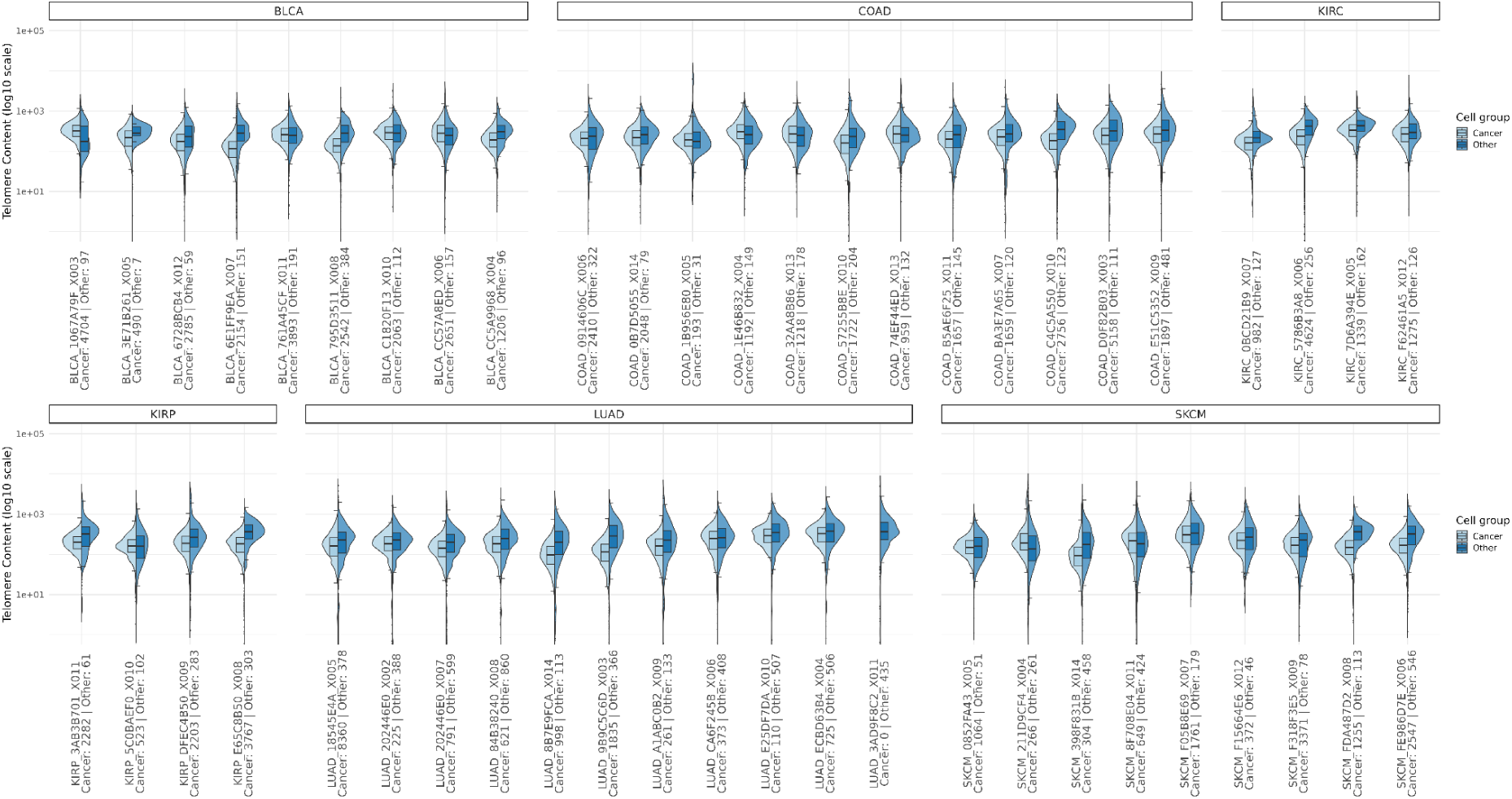
Telomere content distributions across additional TCGA cohorts. Split violin plots of telomere content (y-axis, log10 scale) per patient for BLCA, COAD, KIRC (top), and KIRP, LUAD, SKCM (bottom). Distributions are shown separately for cancer and other cell types (microglia, B cells, T cells, plasma cells, macrophages, endothelial cells, and fibroblasts), with sample labels annotated with the respective cell counts.

**Figure S7.**
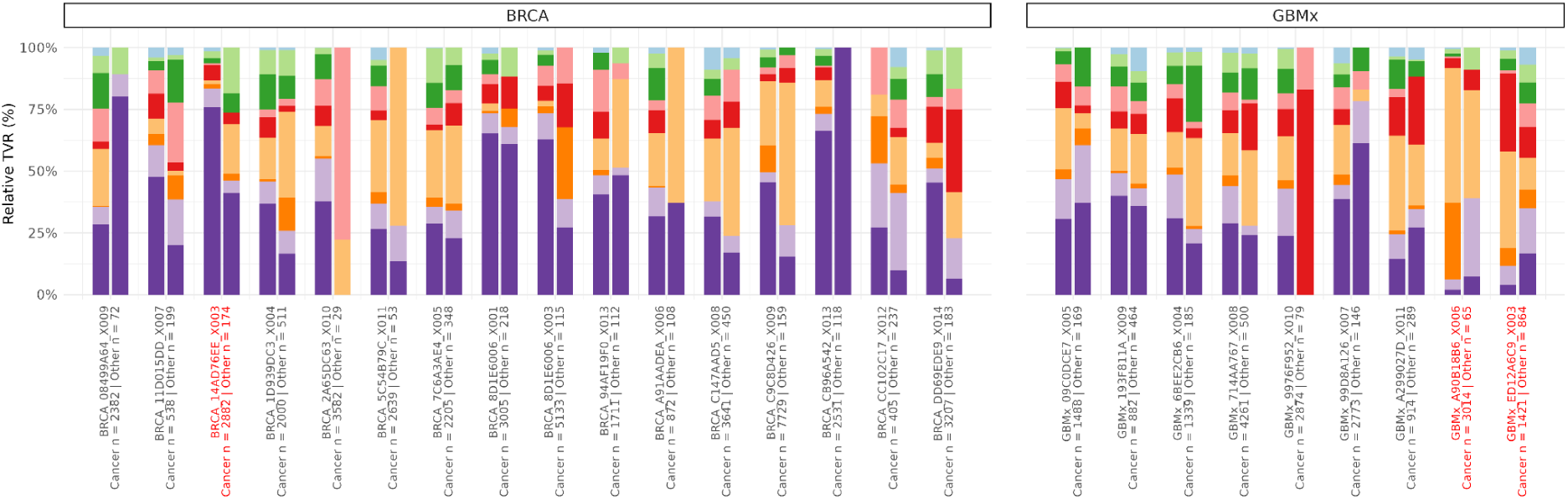
Relative sTVR composition per patient. For each BRCA and GBMx patient and compartment (cancer cells vs. others consisting of microglia, B cells, T cells, plasma cells, macrophages, endothelial cells, and fibroblasts), sTVR counts were summed by TVR class and then normalized to the total sTVR signal within that compartment. The y-axis shows percent of total sTVR signal.

**Figure S8.**
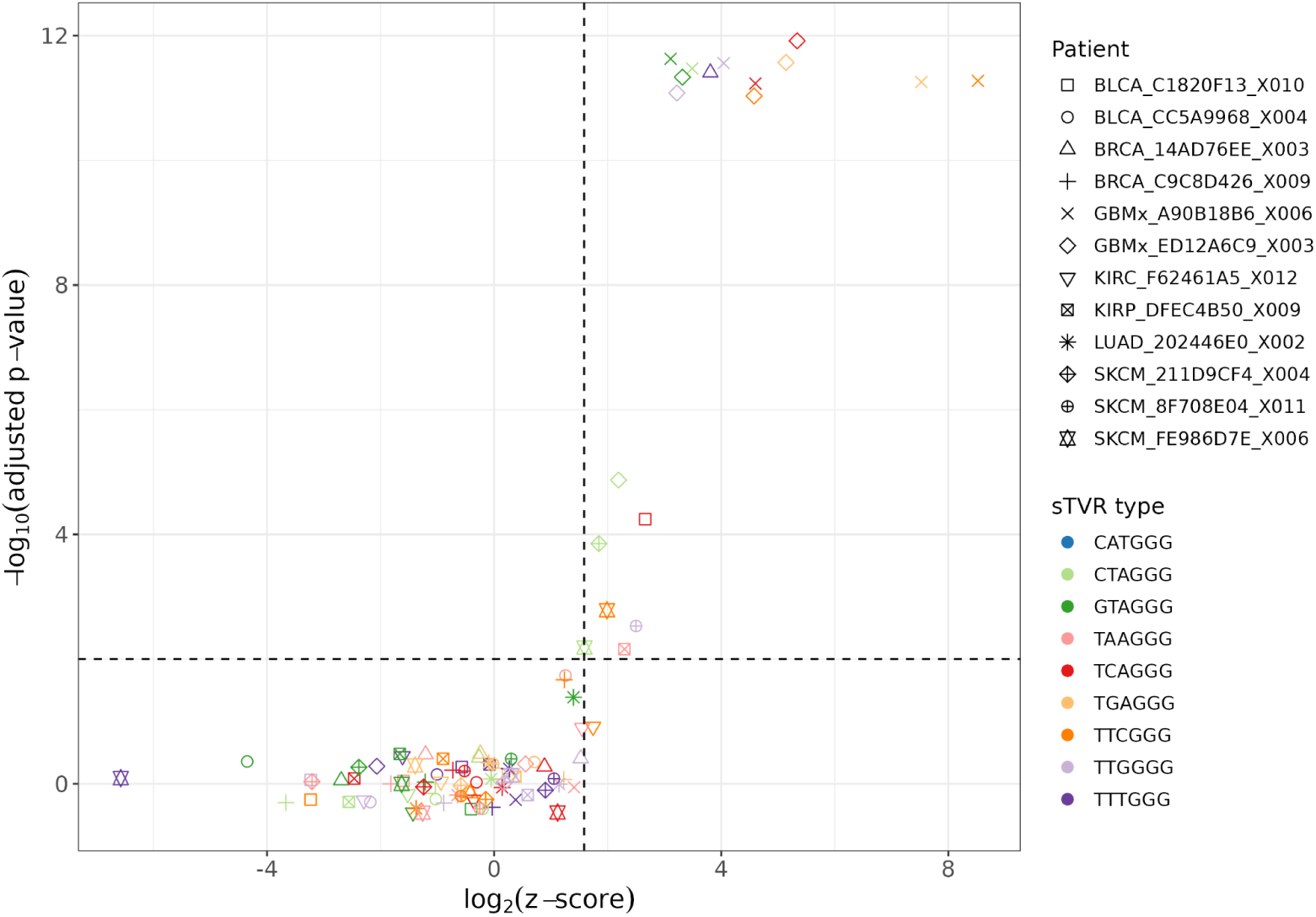
Volcano plot of patient-specific singleton telomere variant repeat derivations in tumor cells. Displayed are all patients exhibiting at least one sTVR type with an absolute z-score greater than three. Each point represents a specific sTVR type per patient, color-coded by TVR sequence, with point shape distinguishing individual patients. The x-axis shows the log2-transformed absolute z-scores, and the y-axis displays the log10 of the Benjamini-Hochberg adjusted p-values. Dashed lines indicate the z-score threshold (|z| = 3) and the significance cut off (-log10(q) = 2). Complete results, including z-scores and adjusted p-values for these patients are provided in Table S3.

**Figure S9.**
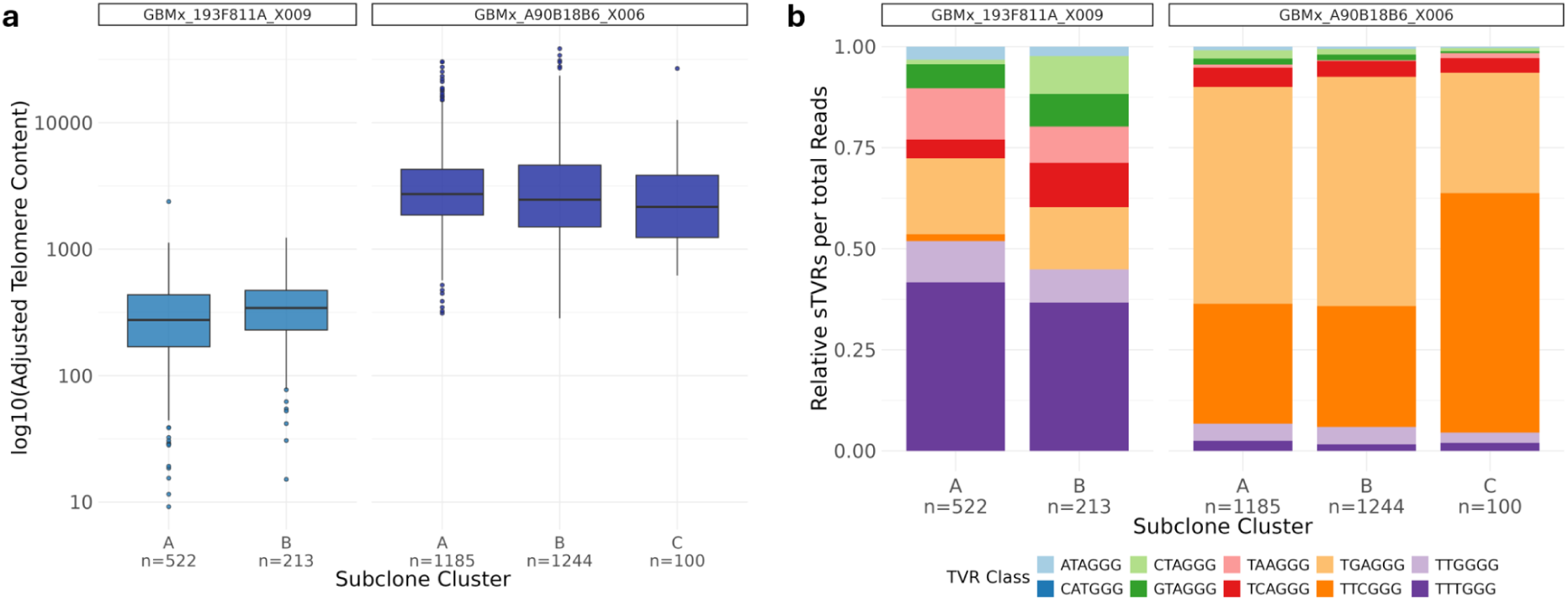
Telomere content and sTVR composition across subclones in selected GBMx patients. **(a)** Boxplots of telomere content (log-scaled openness adjusted telomere content, y-axis) across subclone clusters for two GBMx patients. Each cluster is annotated with the corresponding number of cells (n). **(b)** Relative sTVR composition within the same subclone clusters. Stacked bars show the proportional contribution of individual TVR classes to the overall telomeric signal.

**Table S4.** R packages used in this study. List of R packages used for data analysis in this study.

| Package | Version |
| --- | --- |
| ArchR | 1.0.3 |
| BiocManager | 1.30.25 |
| cowplot | 1.1.3 |
| dplyr | 1.1.4 |
| forcats | 1.0.0 |
| gghalves | 0.1.4 |
| ggplot2 | 4.0.1 |
| ggpubr | 0.6.1 |
| ggrepel | 0.9.6 |
| ggtext | 0.1.2 |
| grid | 4.4.3 |
| gtable | 0.3.6 |
| halfmoon | 0.1.0 |
| purrr | 1.2.1 |
| readr | 2.1.5 |
| rstatix | 0.7.2 |
| reshape2 | 1.4.5 |
| RColorBrewer | 1.1.3 |
| scales | 1.4.0 |
| Seurat | 5.2.1 |
| SingleCellExperiment | - |
| stringr | 1.6.0 |
| tidyr | 1.3.1 |
| viridis | 0.6.5 |

## Notes

### Competing Interest Statement

The authors have declared no competing interest.

### Summary of Updates

This version of the manuscript extends the previous work by incorporating a cell cycle-based correction of telomere content to improve telomere content estimation in single-cell ATAC-seq data, as well as amplification bias-based read filtering. Additionally, the TCGA scATAC-seq atlas was included as an independent dataset for analysis.

